# An integrative somatic mutation analysis to identify pathways linked with survival outcomes across 19 cancer types

**DOI:** 10.1101/017582

**Authors:** Sunho Park, Seung-Jun Kim, Donghyeon Yu, Samuel Pena-Llopis, Jianjiong Gao, Jin Suk Park, Hao Tang, Beibei Chen, Jiwoong Kim, Jessie Norris, Xinlei Wang, Min Chen, Minsoo Kim, Jeongsik Yong, Zabi WarDak, Kevin S Choe, Michael Story, Timothy K. Starr, Jaeho Cheong, Tae Hyun Hwang

## Abstract

Identification of altered pathways that are clinically relevant across human cancers is a key challenge in cancer genomics. We developed a network-based algorithm to integrate somatic mutation data with gene networks and pathways, in order to identify pathways altered by somatic mutations across cancers. We applied our approach to The Cancer Genome Atlas (TCGA) dataset of somatic mutations in 4,790 cancer patients with 19 different types of malignancies. Our analysis identified cancer-type-specific altered pathways enriched with known cancer-relevant genes and drug targets. Consensus clustering using gene expression datasets that included 4,870 patients from TCGA and multiple independent cohorts confirmed that the altered pathways could be used to stratify patients into subgroups with significantly different clinical outcomes. Of particular significance, certain patient subpopulations with poor prognosis were identified because they had specific altered pathways for which there are available targeted therapies. These findings could be used to tailor and intensify therapy in these patients, for whom current therapy is suboptimal.

## Background

In the last few years, studies using high-throughput technologies have highlighted the fact that the development and progression of cancer hinges on somatic alterations. These somatic alterations may disrupt gene function, such as activating oncogenes or inactivating tumor suppressor genes, and thus, dysregulate critical pathways contributing to tumorigenesis. Therefore, precise identification and understanding of disrupted pathways may provide insights into therapeutic strategies and the development of novel agents. Many large-scale cancer genomics studies, such as The Cancer Genome Atlas (TCGA) and the International Cancer Genome Consortium (ICGC), have performed an integrated analysis to draft an overview of somatic alterations in the cancer genome^1–4^. Many of these studies have reported novel candidate cancer genes mutated at high and intermediate frequencies in a specific cancer as well as across many cancer types^4^. However, it is a still challenge to translate somatic mutations in tumors into the pathway model to accurately predict patient clinical outcomes ^5, 6^. Recently, in order to improve the clinical relevance and utility of somatic mutation analyses, Hopfree et al^7^ proposed integrating somatic mutation data with molecular interaction networks for patient stratification. They demonstrated that inclusion of prior knowledge, captured in molecular interaction networks, could improve identification of patient subgroups with significantly different histological, pathological or clinical outcomes and discover novel cancer-related pathways or subnetworks. In a similar manner, other network-based methods have demonstrated that incorporating molecular networks and/or biological pathways can improve accuracy in identifying cancer-related pathways^8–11^.

One limitation of these network-based methods is that they are not designed to fully utilize large-scale somatic mutation data from multiple cancer types to determine which particular pathways are altered by somatic mutations across a range of human cancers. In addition, due to the incomplete knowledge of existing gene set and/or pathway database, these methods are limited to detect pathways based on a number of altered genes annotated in existing gene set and pathway databases. Alternatively, the methods that build pathways *de novo* without incorporating biological prior knowledge can be applicable to detect altered pathways, but these methods were not designed to detect cancer-type specific or commonly altered pathways either.

To address these, we developed an algorithm named NTriPath (Network regularized non-negative TRI matrix factorization for PATHway identification) to integrate somatic mutation, gene-gene interaction networks and gene set or pathway databases to discover pathways altered by somatic mutations in 4,790 cancer patients with 19 different types of cancers. Incorporating existing gene set or pathway databases enables NTriPath to report a list of altered pathways across cancers, and make easy to determine/compare which particular pathways are altered in a particular cancer type(s). In particular, the use of large-scale genome-wide somatic mutations from 4,790 cancer patients and gene-gene interaction networks enables NTriPath to classify genes, which were not annotated in existing gene set or pathway databases, as new member genes of the identified altered pathways based on modular structures of mutational data within a cancer type and/or across multiple cancer types (using matrix factorization) and connectivity in the gene-gene interaction networks. The questions that we investigate here are: first, whether large-scale integrative somatic mutation analysis can reliably identify cancer-type-specific or commonly altered pathways by somatic mutations across cancers; second, whether the identified pathways can be used as a prognostic biomarker for patient stratification - with the assumption that the altered pathways contribute to cancer development and progression and, thus, impact survival.

In these experiments, we demonstrated that the cancer-type-specific and commonly altered pathways identified by NTriPath are biologically relevant to the corresponding cancer type and are associated with patient survival outcomes. We also showed that cancer-specific altered pathways are enriched with many known cancer-relevant genes and targets of available drugs including those already FDA-approved. These results imply that the cancer-specific altered pathways can guide therapeutic strategy to target the altered pathways that are pivotal in each cancer type.

## Results

### NTriPath: An integrative somatic mutation analysis for discovering pathways altered by somatic mutations across multiple cancer types

NTriPath integrates somatic mutations with gene-gene interaction networks and a pathway database to discover altered pathways across human cancers. We collected somatic mutation data from TCGA for 4,790 patients and 19 different cancer types (Table 1). A diagrammatic description of our algorithm is depicted in Figure 1. Four types of data were used as input for our algorithm. First, we generated a binary matrix (***X***) of patients x genes, with ‘1’ indicating a mutation and ‘0’ no mutation. Second, we constructed gene-gene interaction networks (***A***). Third, we incorporated a pathway database (***V*_0_**) (e.g., conserved 4,620 subnetworks across species^12^). Fourth, we included clinical data on the patient’s tumor type (***U***). NTriPath produces two matrices as output; 1) altered pathways by mutated genes (***V***) and 2) altered pathways by cancer type matrix (***S***). The use of both large-scale somatic mutation profiles and gene-gene interaction networks enabled NTriPath to identify cancer-related pathways containing known cancer genes mutated at different frequencies across cancers with newly added member genes according to high network connectivity (***V***) ^8, 13^. Finally we use the altered pathways by cancer type matrix (***S***) to identify altered pathways that are specific for each cancer type. For further details, please see the Materials and Methods section. Our method is also available at www.taehyunlab.org.

**Figure 1.**
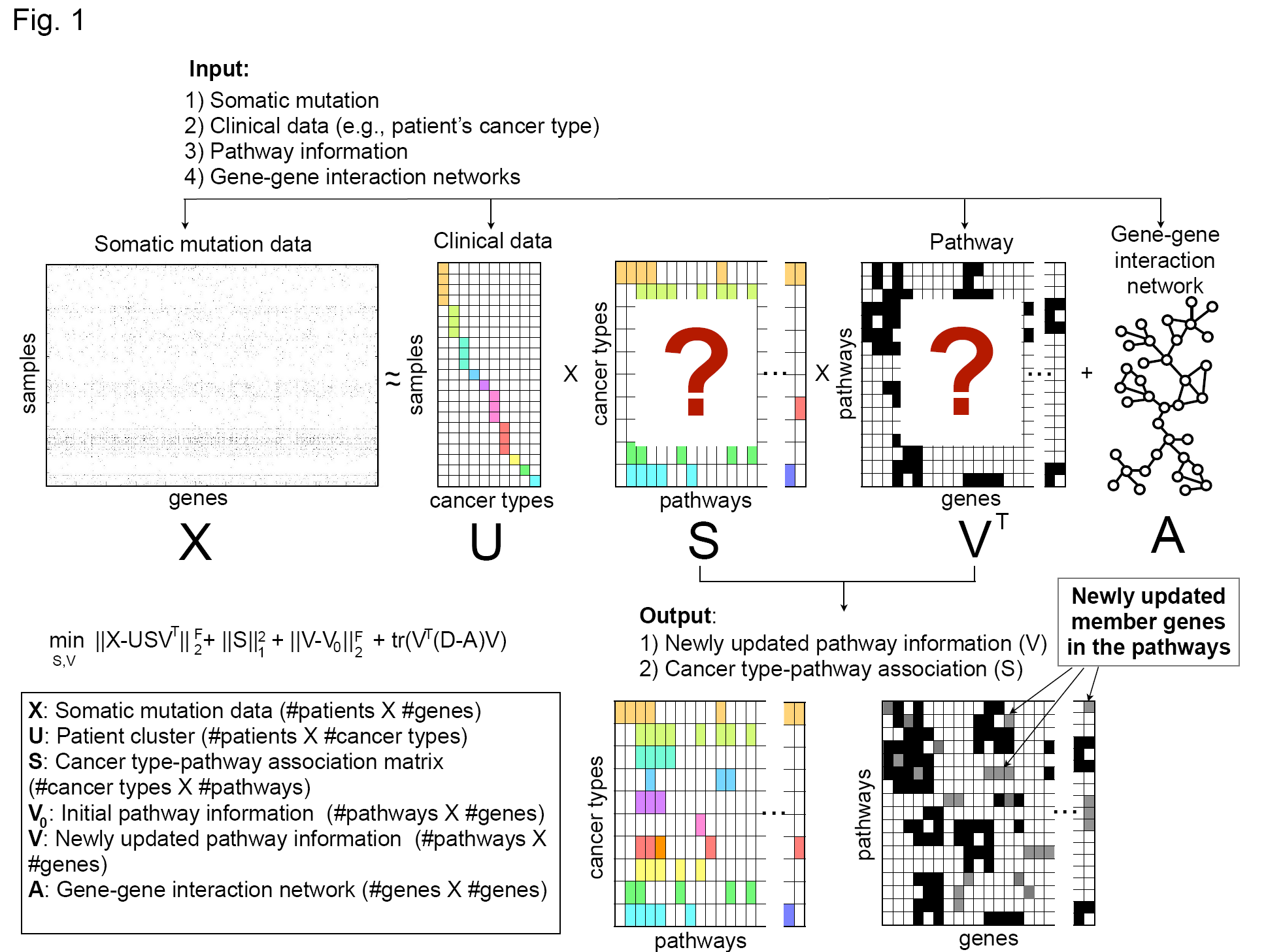
Overview of NTriPath. This figure describes steps to discover altered pathways across multiple cancer types. The aim of the approach is to integrate the somatic mutation data with gene-gene interaction networks and a pathway database for discovering altered pathways across cancers.

**Table 1.** A list of cancer types used in the analysis.

|  | Cancer Type | Number of Patients |
| --- | --- | --- |
| 1 | Acute Myeloid Leukemia (LAML) | 75 |
| 2 | Bladder Urothelial Carcinoma (BLCA) | 136 |
| 3 | Brain Lower Grade Glioma (LGG), | 217 |
| 4 | Breast Invasive Carcinoma (BRCA) | 772 |
| 5 | Cervical Squamous Cell Carcinoma and Endocervical Adenocarcinoma (CESC) | 41 |
| 6 | Colon Adenocarcinoma (COAD) | 270 |
| 7 | Glioblastoma Multiforme (GBM) | 290 |
| 8 | Head and Neck Squamous Cell Carcinoma (HNSC) | 323 |
| 9 | Kidney Chromophobe Renal Cell Carcinoma (KICH) | 65 |
| 10 | Kidney Renal Clear Cell Carcinoma (KIRC) | 210 |
| 11 | Kidney Renal Papillary Cell Carcinoma (KIRP) | 111 |
| 12 | Lung Adenocarcinoma (LUAD) | 380 |
| 13 | Ovarian Serous Cystadenocarcinoma (OV) | 463 |
| 14 | Prostate Adenocarcinoma (PRAD) | 171 |
| 15 | Rectum Adenocarcinoma (READ) | 116 |
| 16 | Skin Cutaneous Melanoma (SKCM) | 269 |
| 17 | Stomach Adenocarcinoma (STAD) | 264 |
| 18 | Thyroid Carcinoma (THCA) | 369 |
| 19 | Uterine Corpus Endometrioid Carcinoma | 248 |

### NTriPath identifies cancer-type-specific altered pathways that are biologically and clinically relevant

In each cancer type, we selected the top 3 ranked altered pathways by statistical significance from NTriPath with the 4,620 subnetwork modules to generate cancer-type-specific altered pathways (See Material and Method section and Supplementary Table 1). Interestingly, NTriPath was able to find altered pathways containing not only genes that were frequently mutated but also genes that were mutated in a small subset of patients in each cancer type (Supplementary Table 2 and Supplementary Figure 1). Gene set enrichment analysis using the genes from the top 3 altered pathways showed that the altered pathways are significantly enriched with well-known cancer-related genes from COSMIC database ^14^ and known drug target genes as well as cancer-relevant biological processes (Supplementary Table 3 and 4).

Focusing on kidney renal clear cell carcinoma (KIRC) as a proof of concept, NTriPath identified the pathway consisting of *VHL, USP33, DIO2*, *TCEB1* and *TCEB2* as the top-ranked altered pathway in KIRC (Figure 2A). The *VHL* (*von-Hippel Lindau*) gene is a well-known tumor suppressor associated with KIRC, and is frequently mutated in patients with KIRC^15–18^. *VHL* was the most frequently mutated gene in TCGA KIRC with 55.7% of patients harboring mutations in the gene. *TCEB1* is mutated at very low frequency in TCGA KIRC cohort. A recent study found that *TCEB1* is mutated in about 3% of the KIRC patients without *VHL* inactivation, and found *TCEB1* preventing the binding of Elongin C to *VHL,* which inactivates the *VHL* pathway^16^. The second highest ranked pathway contained *EP300* and *TP53*. *EP300* and *TP53* were mutated in 8.1% and 5.2% of patients, respectively. *EP300* has been identified as a co-activator of hypoxia-inducible factor 1 alpha (*HIF1α*, whose activation is a hallmark of KIRC tumors. *TP53* was previously found to be associated with poor outcome in TCGA KIRC^19^. The third highest ranked pathway contains *LRP1* and matrix metalloproteinases (*MMP1*, *MMP7*, *MMP9*, *MMP26*). *LRP1* is mutated in 10% of TCGA KIRC cohort, but matrix metalloproteinases (*MMPs*) were not mutated in TCGA KIRC cohort. Biological and clinical relevance of *LRP1* mutation in KIRC has not been previously reported. *MMPs* have been implicated in different types of cancer progression including the acquisition of invasive and metastatic properties in many cancer types. The aberrant expression of *MMPs* has been associated with poor patient survival and prognosis in KIRC patients ^8, 20^. Interestingly, recent studies suggested that *LRP1* induces the expression of matrix metalloproteinase (MMPs) and thus promotes cancer cell invasion and metastasis in many cancers including KIRC^16,21–23^.

**Figure 2.**
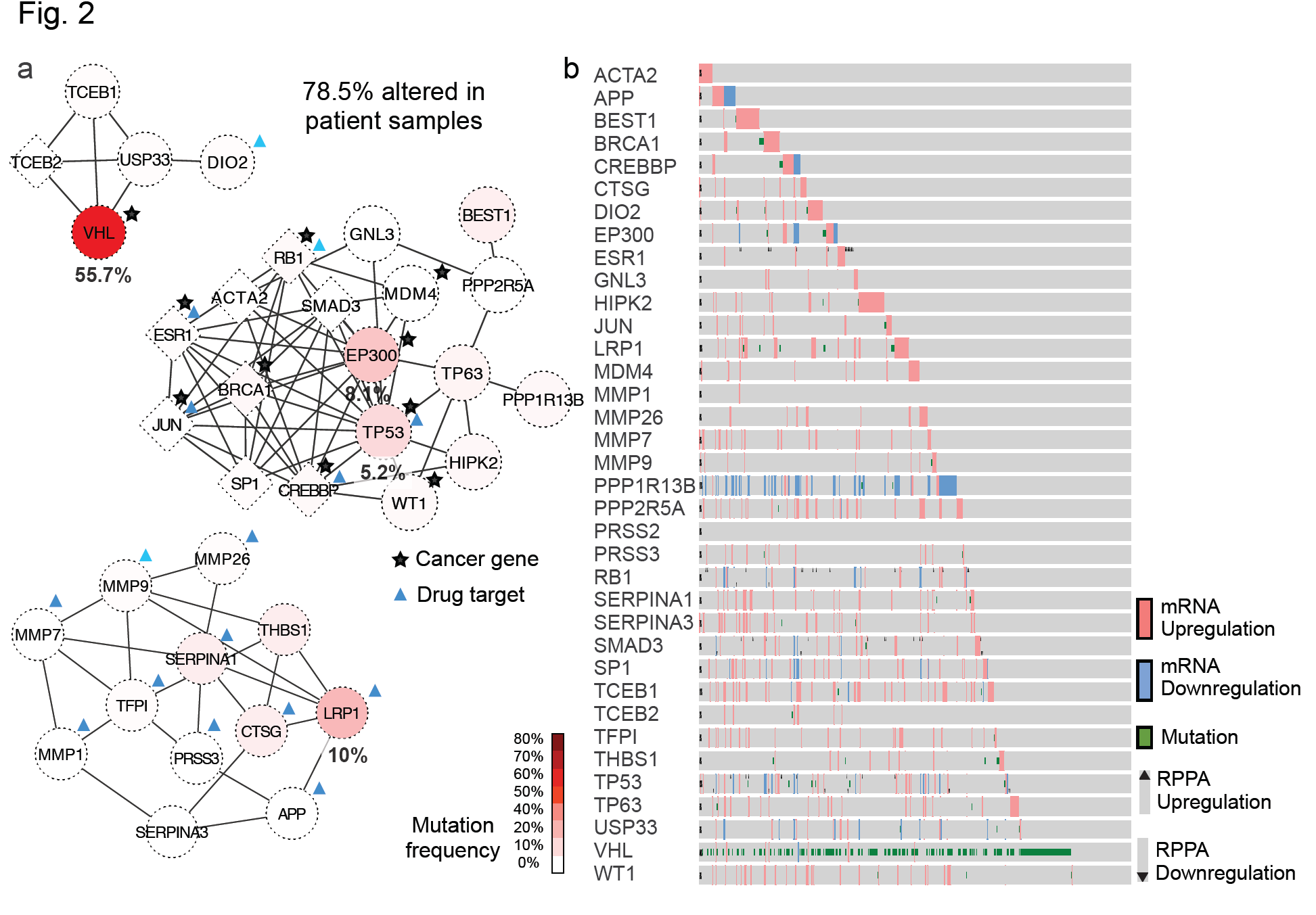
KIRC-specific altered pathways. (A) Diagrams of the top three ranked altered pathways in patients with KIRC. Red color indicates genes that are frequently mutated. A circular-shaped node represents the original member genes annotated in the pathway database, and a diamond-shaped node represents newly identified members genes of the pathways by NTriPath (B) Protein and mRNA expression and mutation status for all genes identified in the top three KIRC altered pathways. Each row represents a member gene in the TCGA KIRC-specific altered pathway, and each column represents a patient sample in TCGA KIRC cohort.

NTriPath identified many new member genes in the top ranked pathways including *TCEB2*, *JUN*, and *SP1* as well as other tumor suppressors such as *CREBBP*, *SMAD3*, *BRCA1* and *RB1*. These newly identified member genes by NTriPath were mutated at a very low frequency or not mutated at all in TCGA KIRC patients. Instead, these genes interacted with many frequently mutated genes in the networks and were often dysregulated at the mRNA and protein levels in many KIRC patients (Figure 2B). For example, *TCEB2*, *SP1* and *JUN* were not mutated but yet their expression was dysregulated in 7%, 10% and 2% of TCGA KIRC patients, respectively. Previous studies have shown that dysregulation in *TCEB2* is expected to disrupt the protein complex that ubiquitinates *HIF1α,* resulting in the same phenotype as *VHL* inactivation by mutation or promoter hypermethylation^24–26^. In addition, *SP1* and *JUN* were previously identified as major transcriptional regulators associated with signaling circuit to promote tumor growth and invasion in KIRC ^18^.

Taken together, these results demonstrate that NTriPath is an effective tool to accurately identify cancer-specific altered pathways including known cancer genes mutated at a high or intermediate frequency in the patients, as well as genes mutated at a very low frequency or not mutated at all yet may be fundamental role in development and/or progression of KIRC.

### Cancer-type-specific altered pathways across cancer types correlate with patient survival outcomes

We hypothesized that cancer-type-specific altered pathways reflect the molecular basis underlying the patient clinical outcomes. This would allow us to use member genes in the altered pathways as gene signatures to stratify patients into subgroups with different clinical outcomes for each type of cancer. We first collected a dataset consisting of gene expression profiles from 3,656 patients with their survival information from TCGA cohorts. We then used member genes in the top 3 ranked cancer-type-specific altered pathways to perform consensus clustering for each cancer type (see Material and Method section). We generated Kaplan-Meier (KM) curves based on the groups produced by consensus clustering and found that patient survival was significantly different among the groups (Figure 3 and Supplementary Figure 2). In TCGA KIRC, we found three patient subgroups (A, B and C), with the Group C having the poorest survival. A log-rank test indicated that Groups A and C had significantly different survival outcomes (Log-rank test p-value = 1.840e-08, Hazard ratio = 2.94) with median survival times of 41.9 months for group A compared to 30.8 months for group C (Figure 3A). Other examples are Bladder Urothelial Carcinoma (BLCA), Head and Neck squamous cell carcinoma (HNSC), and Skin Cutaneous Melanoma (SKCM) patient subgroups identified by NTriPath pathway signatures. While the molecular classification of clinically relevant subtypes of these cancers is still challenging, we found patient subgroups having significantly different survival in these cancers (Log-rank test p-value= 0.0086, 0.0010, and 0.0210, repectively) (Figure 3B, 3C and 3D).

**Figure 3.**
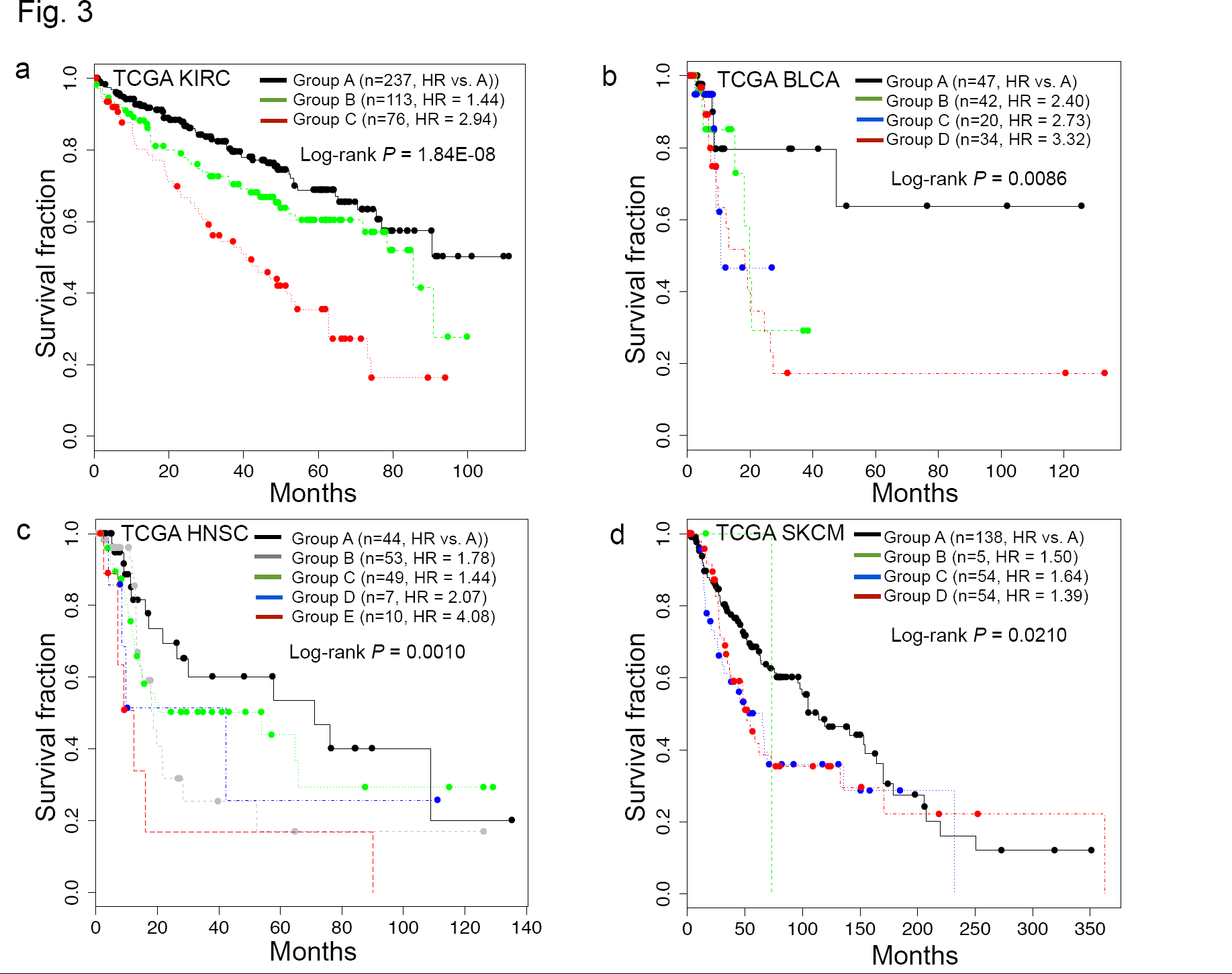
Cancer-type-specific altered pathways across cancers correlate with survival outcomes. Kaplan-Meier survival plots based on patient subgroups defined by consensus clustering using genes from the top 3 altered pathways for (a) kidney renal cell carcinoma (KIRC), (b) bladder urothelial carcinoma (BLCA), (c) head and neck squamous carcinoma (HNSC), and (d) skin cutaneous melanoma (SKCM).

Experiments with other TCGA datasets, including those for Breast invasive carcinoma (BRCA), Glioblastoma Multiforme (GBM), Lung adenocarcinoma (LUAD), and Ovarian serous cystadenocarcinoma (OV) consistently showed that the use of member genes in cancer-type-specific altered pathways could serve as a prognostic biomarker for patient stratification (Supplementary Figure 2 and 3). For comparison, we also attempted to cluster patients using significant frequently mutated genes previously identified by the TCGA Pan-Cancer study^1^. The results of consensus clustering using the NTriPath-derived pathway signatures and the TCGA Pan-Cancer-derived mutated gene signatures showed that the results from NTriPath-derived pathway signatures had higher significance levels for BLAC, BRCA, and KIRC, and comparable results for the GBM, HNSC, and LUAD cancer types (Figure 4). These findings suggested that NTriPath-derived altered pathways could be used as prognostic biomarkers for better patient stratification.

**Figure 4.**
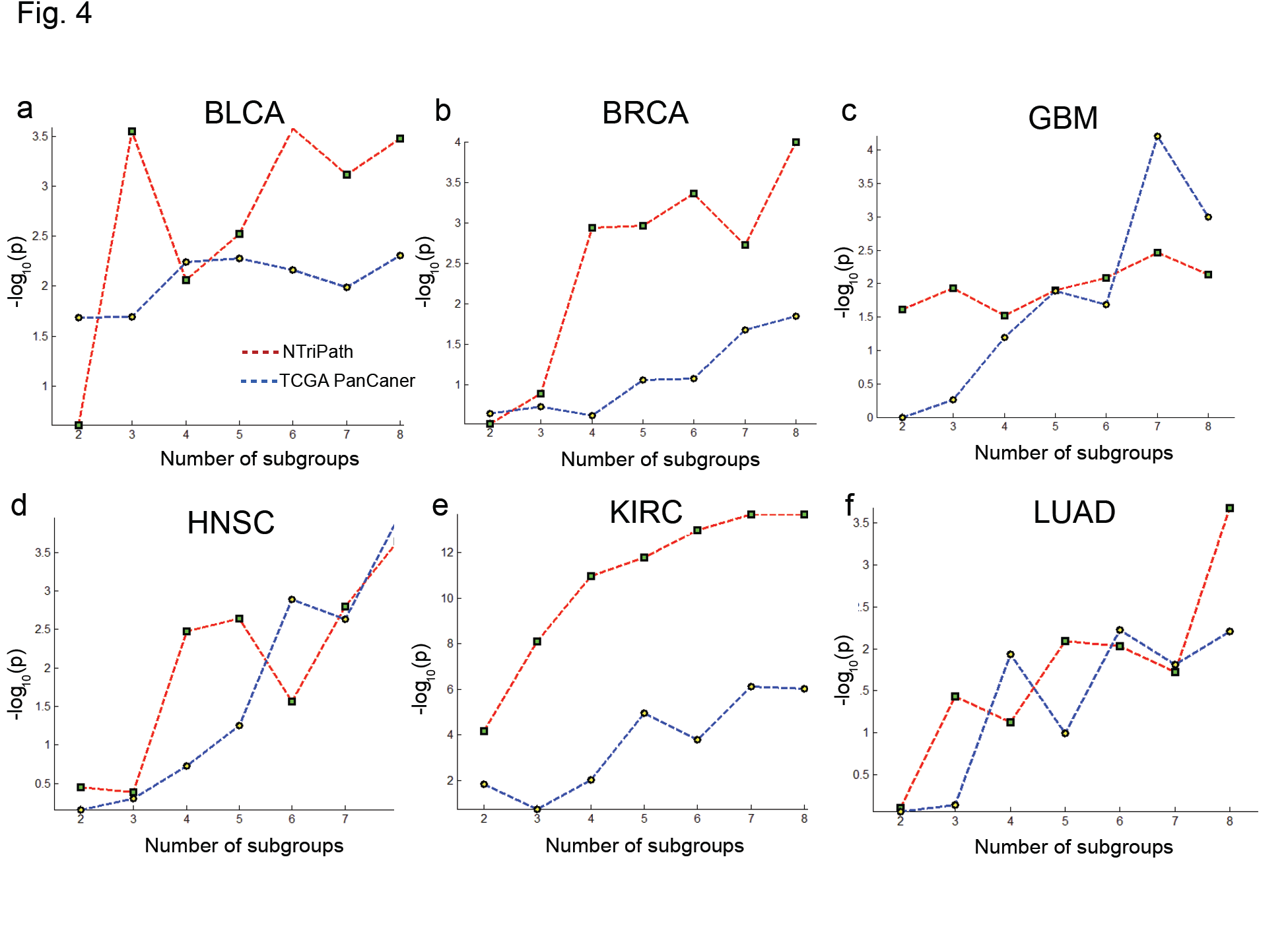
Comparing NTriPath-derived signatures with mutation-frequency-based signatures. This figure describes comparisons of patient stratification using signatures derived from NTriPath and mutation frequency reported in Kandoth, C. *et al.* ^1^.

### Independent cohorts for the validation of the cancer-type-specific altered pathways

We performed multiple validations to evaluate the robustness and the reproducibility of NTriPath. First, we evaluated the robustness of the cancer-type-specific altered pathways identified in the TCGA cohort for prognostic stratification. We generated gene expression profiles of 102 HNSC patients from our institution and used the member genes of the top 3 HNSC cancer-type-specific altered pathways in the TCGA cohort for patient stratification. In addition, we also used publically available gene expression data from two ovarian cancer datasets, one lung cancer dataset, two colon cancer datatsets for a total of 1,112 patients, and used the top 3 cancer-type specific altered pathways for corresponding cancer type for independent validation. In the HNSC cohorts, we found six patient subgroups (A through F), with the group F patients having the poorest survival times (Figure 5A). A log-rank test indicated that groups A and F had significantly different survival outcomes (p-value = 0.038, hazard ratio = 1.88) with median survival times of 78.1 months for group A and 26.7 months for group F. Similarly, we found patient subgroups having significantly different survival outcomes in lung cancer, ovarian cancer, and colorectal cancer (Figure 5B-D and Supplementary Figure 3). Secondly, we verified the reproducibility of NTriPath for the identification of the cancer-type-specific altered pathways. We collected the level 2 somatic mutation data from 19 human cancer types those were updated after we collected initial dataset used in the original experiments from the TCGA data portal. We found that there are 1891 newly updated patients’ mutation data from 15 cancer types (see Supplementary Table 5). We re-ran NTriPath to identify cancer-type-specific pathways across 19 cancers using 6681 patients’ somatic mutation data including those of newly updated patients’ mutation data. Interestingly, we found that many top ranked pathways identified by NTriPath in the original experiments were consistently highly ranked in the new experiments (see Supplementary Table 6).

**Figure 5.**
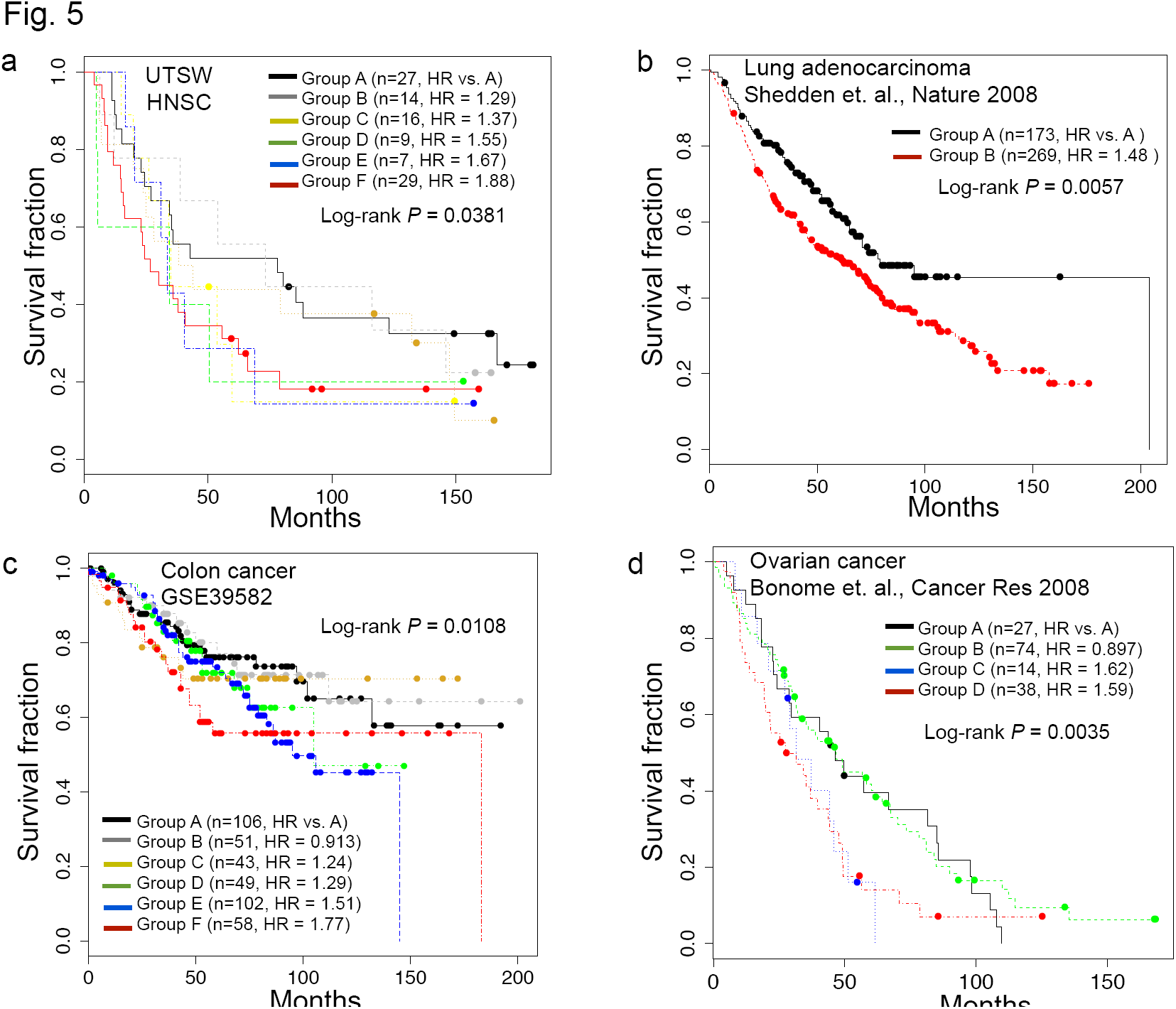
Validation in independent cohort. This figure describes Kaplan-Meier survival plots for patient subgroups from (a) UTSW HNSC (b) Lung adenocarcinoma (c) Colon cancer, (d) Ovarian cancer.

These results reassure that NTriPath is a robust tool to detect the altered pathways across cancers, and the altered pathways identified by NTriPath can serve as robust prognostic signatures for identifying patient subgroups with different survival outcomes across multiple cancer types.

### NTriPath identified potential therapeutic targets in poor prognosis patient subgroups

We further investigated whether we could identify potential targets or the therapy for the identified poor prognosis patient subgroups. Interestingly, we found that many known drug targets in the cancer-type specific altered pathways are often up-regulated in poor prognosis patient subgroups across cancers (see Method section and Supplementary 7). For example, in TCGA KIRC cohort, *LRP1* and *MMP9,* targets of FDA-approved drugs Tenecteplase and Captopril, were significantly up-regulated in poor prognosis group compared to good prognosis group (FDR-adjusted p-value < 0.05 with t-test). Tenecteplase binds to *LRP1* and induces both *LRP1* and *MMP9* expression, and Captoprilare inhibits *MMP9* expression. Thus, combinatorial therapy of these drugs can be beneficial for the KIRC patients with high *LRP1* and *MMP9* expression^27–44^. Another notable example includes DNA Topoisomerase I (*TOP1),* a target of well-known FDA-approved anticancer drugs such as Irinotecan and Topotecan, identified by NTriPath as a new member gene into the cancer-type-specific altered pathways across many cancers including HNSC. Interestingly, we found that *TOP1* was up-regulated in poor prognosis subgroups in HNSC from both TCGA and UTSW cohorts (Group E and F in Figure 3C and 5A, respectively). In addition, we found that some patients with overexpression of *TOP1* in TCGA HNSC poor prognosis subgroup have developed therapy resistance against single chemotherapeutic agent such as Cisplatin. Interestingly, there is an ongoing trial in advanced HNSC showing efficacy of *TOP1* inhibitor Irinotecan with Cisplatin in a poor prognosis patient subgroup ^45^. These observations may suggest that *TOP1* inhibitors-based combinations might offer an effective treatment option for HNSC patients with overexpression of *TOP1*. Taken together, these findings suggested that the use of NTriPath-derived altered pathways containing available drug targets may allow for the development of more tailored therapeutics.

## Discussion

Systematic understanding of how somatic mutations influence clinical outcomes is essential for the development and application of personalized therapies. Especially organizing alterations at the individual gene level and in the molecular pathways can correlate altered pathways and vulnerabilities with specific genetic lesions, and provide novel insights into cancer biology, biomarkers for patient stratification in clinical trials, and potential targeted drug development^46^. Here, we systematically identified biological and clinical relevant cancer-type-specific altered cross multiple cancer types. In particular, the integration of somatic mutation with biological prior knowledge led to the identification of altered pathways that contain recurrently mutated genes as a hallmark of specific cancer types. Interestingly, we found that several genes, while not frequently mutated or not mutated at all in patients, were part of cancer-type-specific altered pathways that have been causally implicated in the development of corresponding cancer types, and expressions of those genes are significantly associated with clinical outcomes (Supplementary Figure 4). For example, no mutation of *MMP7* has been reported, but high expression of *MMP7* (p = 0.00191, HR=1.7 (95%CI,1.21-2.38)) is significantly associated with poor survival in TCGA KIRC patients. Other examples include *CABLES1* (p = 0.00272, HR=0.486 (95%CI,0.301-0.787)) in TCGA HNSC and LUAD, and *GCH1* (p = 0.0000528, HR=0.52 (95%CI,0.367-0.763)) in TCGA SKCM are not frequently or not mutated but low or high expression of those genes are significantly associated with poor survival. In addition, we found that known drug targets are not frequently mutated but often up-regulated in poor prognosis patient subgroups across many cancers. These results further corroborate that the integrative analysis of somatic mutations with additional biological prior knowledge may elucidate potential candidate genes associated with clinical outcomes and could be potentially used to design targeted therapy, which cannot be readily identified by somatic mutation analysis alone.

In our analysis, we did not remove synonymous mutations or further select a shorter list of recurrent mutated genes in cohorts with stringent criteria either ^3, 47^. However, Hopfree et al^7^ showed that filtering synonymous mutations resulted in a decreased ability to detect patient subgroups with different survival outcomes. Another recent study also showed that synonymous mutations could affect functions of oncogene and tumor suppressors ^48^. In addition, our experimental results for patient stratification in comparison with recurrent mutated gene signatures identified by the TCGA Pan-Cancer^1^ indicated that the use of NTriPath-derived pathways showed a comparable or better performance in discovering patient subgroups with different survival outcomes across cancers. To evaluate the impact of different network resources, we used networks from the HPRD ^49^ and Rossin, E.J. *et al*^50^ and repeated experiments. We summarize the results of altered pathways and patient stratification using different network resources and provide on our supplement website.

Lastly, NTriPath is a general computational algorithm and can be applied to other data types such as gene expression, copy number alteration, and methylation to identify altered pathways by different types of genomic aberrations. NTriPath can also be used to find altered pathways across associated with other cancer-related phenotypes (e.g., patient groups having therapy resistance vs. sensitivity, metastatic vs. non-metastatic).

## Conclusions

We have described an integrative somatic mutation analysis for discovering altered pathways in human cancers. NTriPath integrates somatic mutation data and prior biological knowledge from the pathway database and molecular networks to identify significantly altered pathways and their associations with specific cancer types. Specifically, NTriPath effectively utilizes mutation patterns that exist in only a subset of samples (or specific cancer types), thus revealing pathways altered by complex mutation patterns across cancer types. Furthermore, the use of gene-gene interaction networks and the pathway database provides the potential to identify altered pathways enriched with genes harboring mutations at high/intermediate frequencies, as well as those not mutated per se but nevertheless playing critical roles in tumorigenesis in network and pathway contexts. Thus, NTriPath is uniquely suited to provide a global analysis of altered pathways by somatic mutation across cancer types.

We applied NTriPath to somatic mutation data from 19 types of cancers, and discovered cancer-type-specific altered pathways based on these mutations in human cancers. Functional enrichment analysis of cancer-type-specific pathways demonstrated that the identified cancer-type-specific altered pathways are biologically meaningful to each cancer type. It also provided unique pathway views of key biological processes underlying each cancer type. Of particular significance, we identified a patient subgroup with poor survival by cancer-type-specific altered pathway signatures from TCGA cohorts, which in independent cohorts. These results implied the potential utility of cancer-type-specific altered pathway signatures to serve as a guide to tailored treatment in a patient subgroup.

## Materials and Methods

### Somatic mutation, human gene-gene interaction networks, and pathway data

The level 2 somatic mutation data from 19 human cancer types were collected from the TCGA data portal on May 19^th^ 2013^51^. We constructed a gene-gene interaction network by combining networks from Zhang, S. *et al* ^52^, the Human Protein Reference Database (Dec. 2013)^53^ and Rossin, E.J. *et al*^50^..Four sets of pathways were used in the analysis: 1) 4,620 conserved subnetworks from the human gene-gene interaction network^12^, 2) KEGG, 3) Biocarta, and 4) Reactome gene sets from MsigDB (Sept. 2010)^54^.

### Algorithm

The algorithm identifies pathways disrupted by mutated genes. Disrupted pathways are found based on the factorization results from the network regularized nonnegative tri-matrix factorization.

#### 1 Notations

We construct a binary data matrix ***X*** ∈ *R*^*N*×*M*^ from the mutation data, where *N* is the number of patients, *M* is the number of genes and the (*i*, *j*) ^*th*^ element of the matrix ***X***, [***X***]_*i*,*j*_, is ‘1’ if the *i*th patient has a mutation on the *j*th gene, ‘0’ otherwise. We derive the adjacency matrix from the human gene-gene interaction networks and denote it as ***A***, where [***A***]_*ij*_=‘1’ if the *i*th gene is interacting with the *j*th genes in the networks and ‘0’ otherwise. We define the graph Laplacian matrix by ***L*** = ***D*** - ***A***, where each diagonal element in the diagonal matrix ***D*** is given by [***D***]_*ii*_ = **∑**_*ji*_[***A***]_*ij*_. We construct a binary matrix ***U*** ∈ *R*^*N*×*K*_1_^ denoting patient cluster, where *K*_1_ indicates the number of cancer types and [**U**]_*ij*_ = 1 indicates the *i*th patient has *j*th cancer type. We construct a binary matrix ***V***_0_ ∈ *R*^*M*×*K*_2_^ from the specific pathway database denoting pathway information, where *K*_2_ is the number of pathways and [***V***_0_]_*ij*_ = 1 if the *i*th gene is annotated in *j*th pathway as a member in the pathway database, otherwise 0. Since current pathway database annotation is still incomplete, we define a matrix ***V*** ∈ *R*^*M*×*K*_2_^ denoting newly updated pathway information including newly added member genes by NTriPath. We define a matrix ***S*** ∈ *R*^k_1_×k_2_^ denoting cancer type and pathway associations, where each element of [***S***]_*i*, *j*_ represents associations between *i*th cancer type with *j*th pathway. Higher values of elements indicate stronger associations between cancer types and pathways. Since ***V*** and ***S*** are unknown, we need to learn about those matrices during optimization (see below section for details)

#### 2 Network regularized non-negative tri-matrix factorization

The network regularized nonnegative tri-matrix factorization is an extension of Nonnegative Tri Matrix Factorization (NTMF); in this work, somatic mutation data ***X*** is factorized as the products of three element-wise non-negative matrices ***U***, ***S***, and ***V*** denoting patient’s cancer type, cancer-type and pathway associations, and cancer-related pathways, respectively. We here consider a weighted loss function to deal with the sparseness of the data matrix ***X*** (More than 98% of entries are zero). It enables us to focus on an approximation error at nonzero entries, which correspond to mutated genes. In addition, to incorporate the prior knowledge from human gene-gene interaction networks and pathway datasets into factorizations, we enforce constraints on parameters, which involve the graph Laplaican ***L*** and the pathway information ***V***_0_.

All these ideas are accomplished by minimizing the following objective function

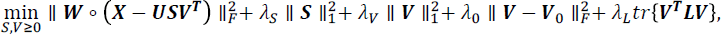

where ***W*** ∈ *R*^*N*×*M*^ is a weight matrix where [***W***]_*ij*_ = 1 if [***X***]_*ij*_ > 0 otherwise 0, and the operator ο represents the element-wise multiplication. Here, we are only interested in learning of ***S*** and ***V*** among three factor matrices, since factor ***U*** can be obtained from the patient’s clinical information.

To solve our minimization problem, we adapt the multiplicative update method for NTMF proposed in a recent study^55^, which contains a routine for avoiding ‘*inadmissible zeros problem’* where the solution of multiplicative update rules is stuck at zero when an entry in the factor becomes.

##### Step 1: Initialization

Initialize the factor matrices ***S*** = **1** and ***V*** = ***V*_0_**, where **1** ∈ *R*^*K*_1_×*K*_2_^ is a matrix whose elements are all one. Set the regularization parameters *λ*_*S*_ = *λ*_*V*_ = *λ*_*L*_ = 1 and *λ*_0_ = 0.1. Set the user specified parameters for avoiding the inadmissible zeros problem, *κ*_*tol*_ = 10^-10^, *κ* = 10^-6^ and *∈* = 10^-10^.

##### Step 2: Iteration

Iterate until it converges or reaches the maximum number of iterations:

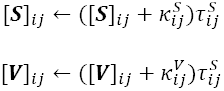

where 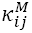 is set to *κ* if [***M***]_*i,j*_ ≥ *κ*_*tol*_ and 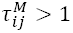, otherwise 0, and

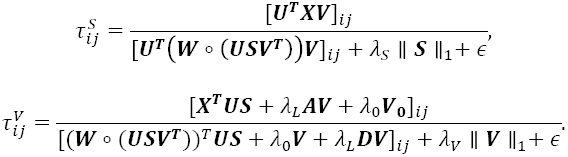

Empirically, the algorithm converges fast within 50 iterations in the experiments.

#### 3 Identification of cancer-type-specific altered pathways

Once the above optimization problem is solved, we used ***S*** matrix to identify cancer-type-specific altered pathways across cancers. Specifically, we ranked pathways based on values of elements of the ***S*** matrix for each cancer type (e.g., rank pathways based on all values of *i*th row which indicate association scores between ith cancer type and all pathways). In addition, to measure statistical significance of cancer-type and pathway associations, we performed a permutation test (e.g., we randomly permuted somatic mutation data and repeated experiments 5,000 times to calculate empirical p-values) and defined cancer-type-specific altered pathways based on the following strict criteria: 1) Pathways must be ranked within the top ***K***th compared to other pathways in each cancer type based on their association scores in matrix ***S***. 2) Pathways must have significant BH-adjusted p-values (Benjamini-Hochberg adjusted p-values using a false discovery rate cutoff of 0.1) (See Supplementary X). In this work, we selected the top 3 ranked pathways having significant BH-adjusted p-values per each cancer type. Top ranked pathways for KICH, KIRP, and THCA were excluded for further analysis, due to the insignificant BH-adjusted p-values.

##### Gene expression data and clustering

We collected RNA-seq data for TCGA BLCA, BRCA, HNSC, KIRC, LAML, LUAD, SKCM, STAD, UCEC from cBioPortal^56–58^ using CGDS MATLAB toolbox with RNA Seq V2 RSEM option. We collected microarray gene expession profiles for TCGA GBM from the TCGA dataportal^51^ and TCGA OV and two others from Zhang, W. *et al*. ^59^. We collected colon cancer data from GSE39582 and lung cancer data from Shedden, K. *et al.* ^60^. RNA-seq data were z-transformed while other expression data were quantile normalized, log transformed, and expression values were median centered. To perform consensus clustering, we used Matlab K-means clustering and used two-way hierarchical clustering.

## Authors’ contributions

SP and THH designed the study. SP and THH performed the analysis. THH supervised the project. SP, JN, SL, and THH wrote the paper. All authors read and approved the final manuscript.

## Acknowledgements

We would like to thank to Quantitative Biomedical Research Center to allow us to use their computational resources.

---

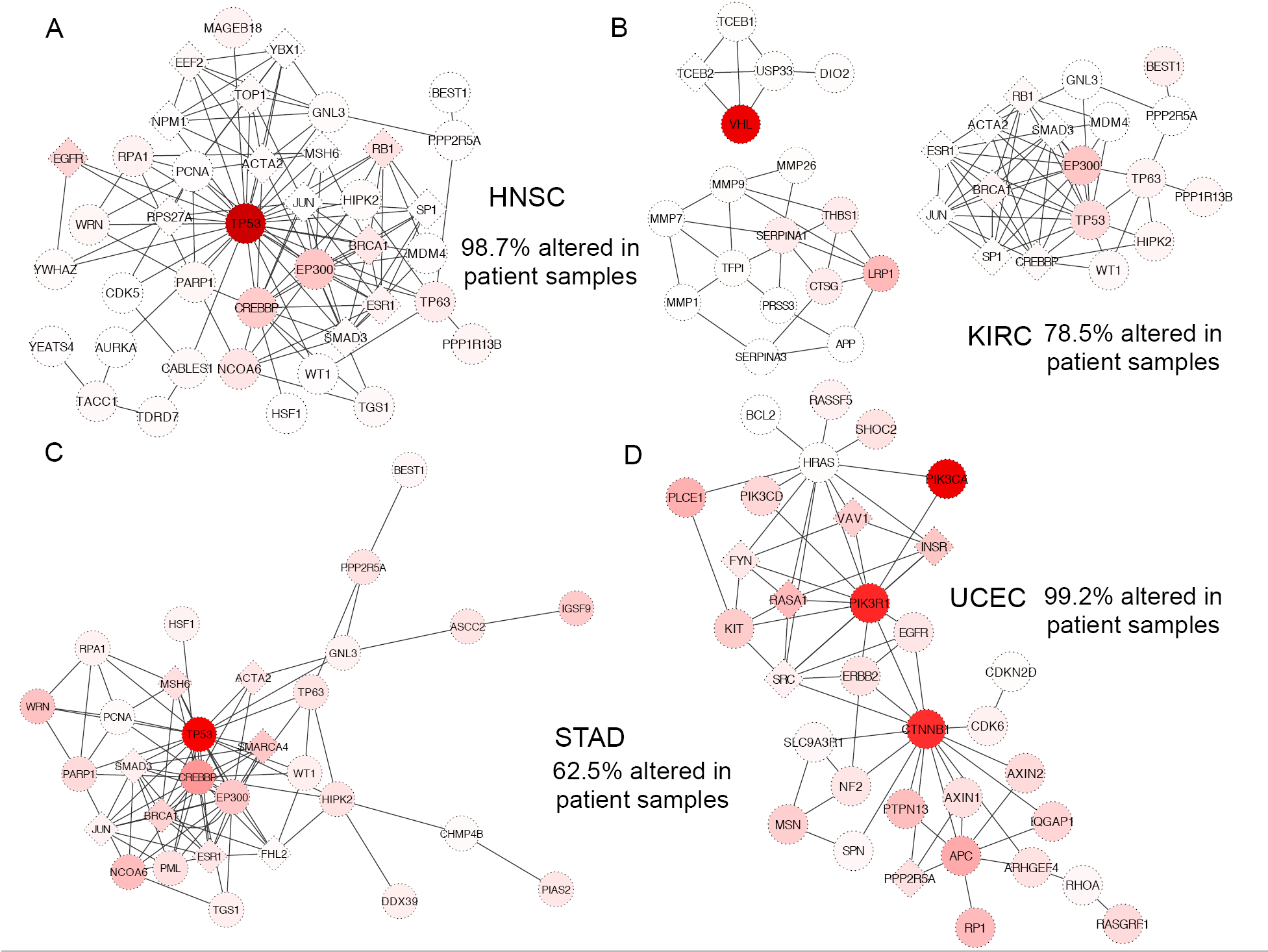
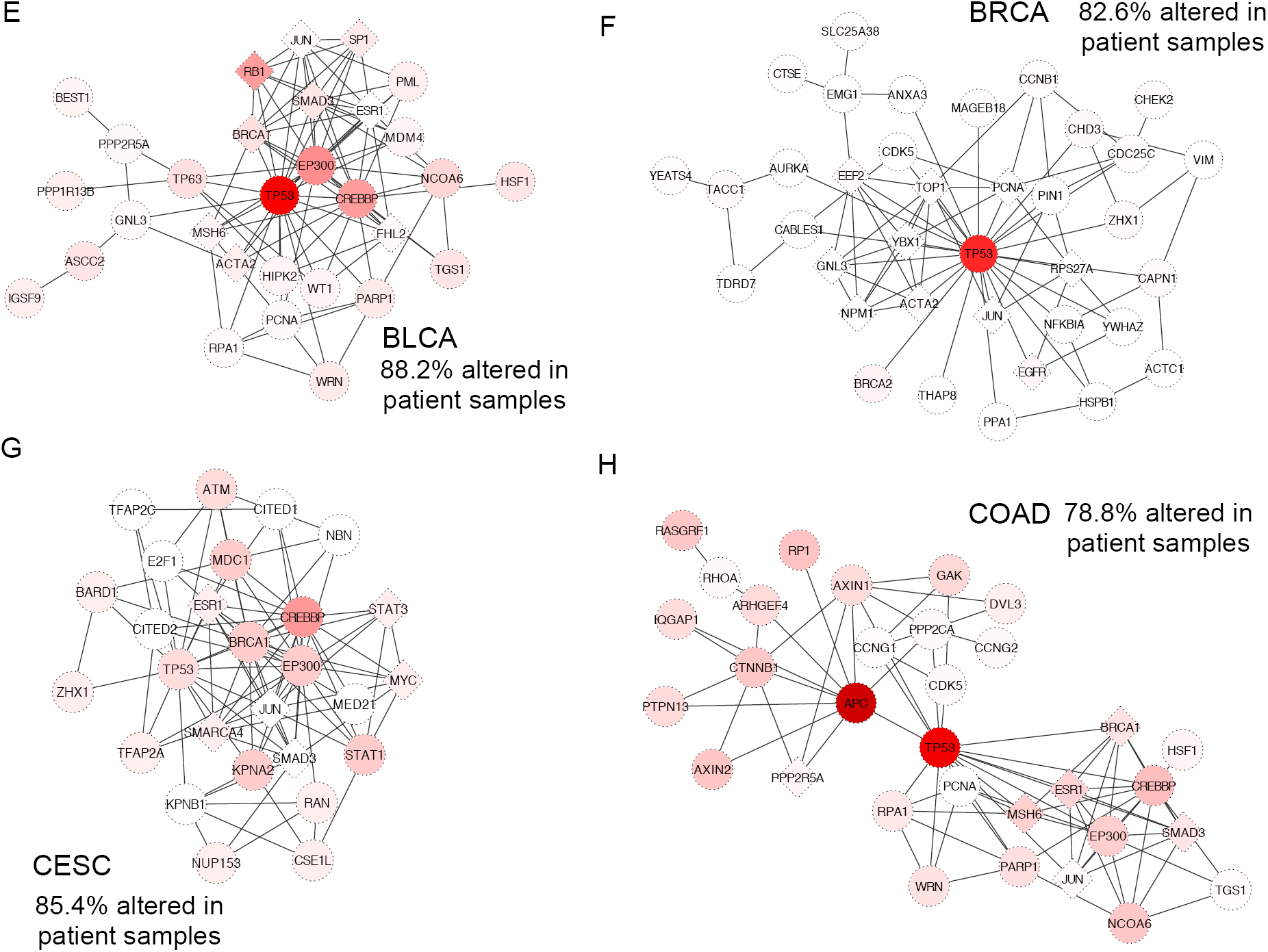
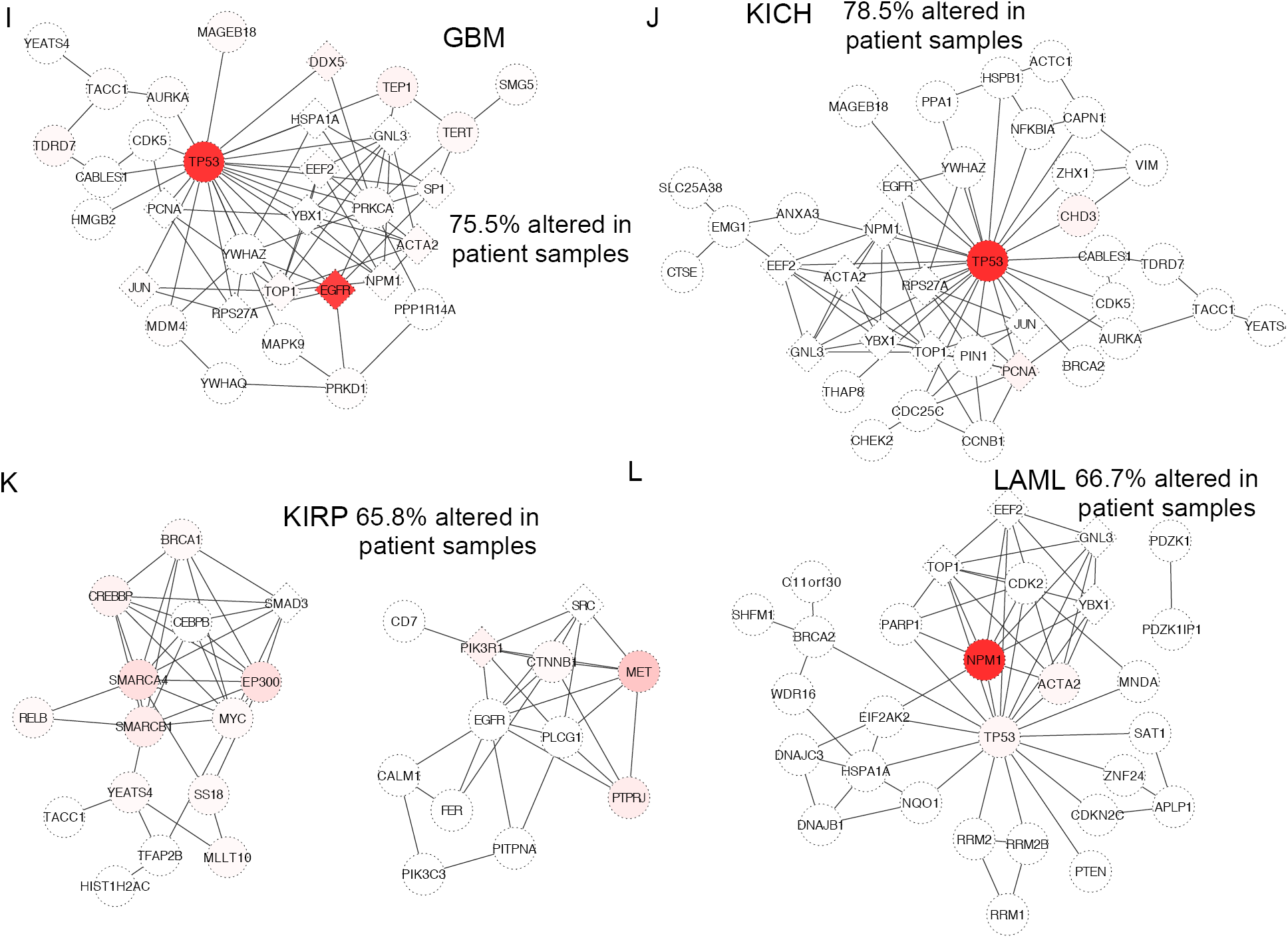
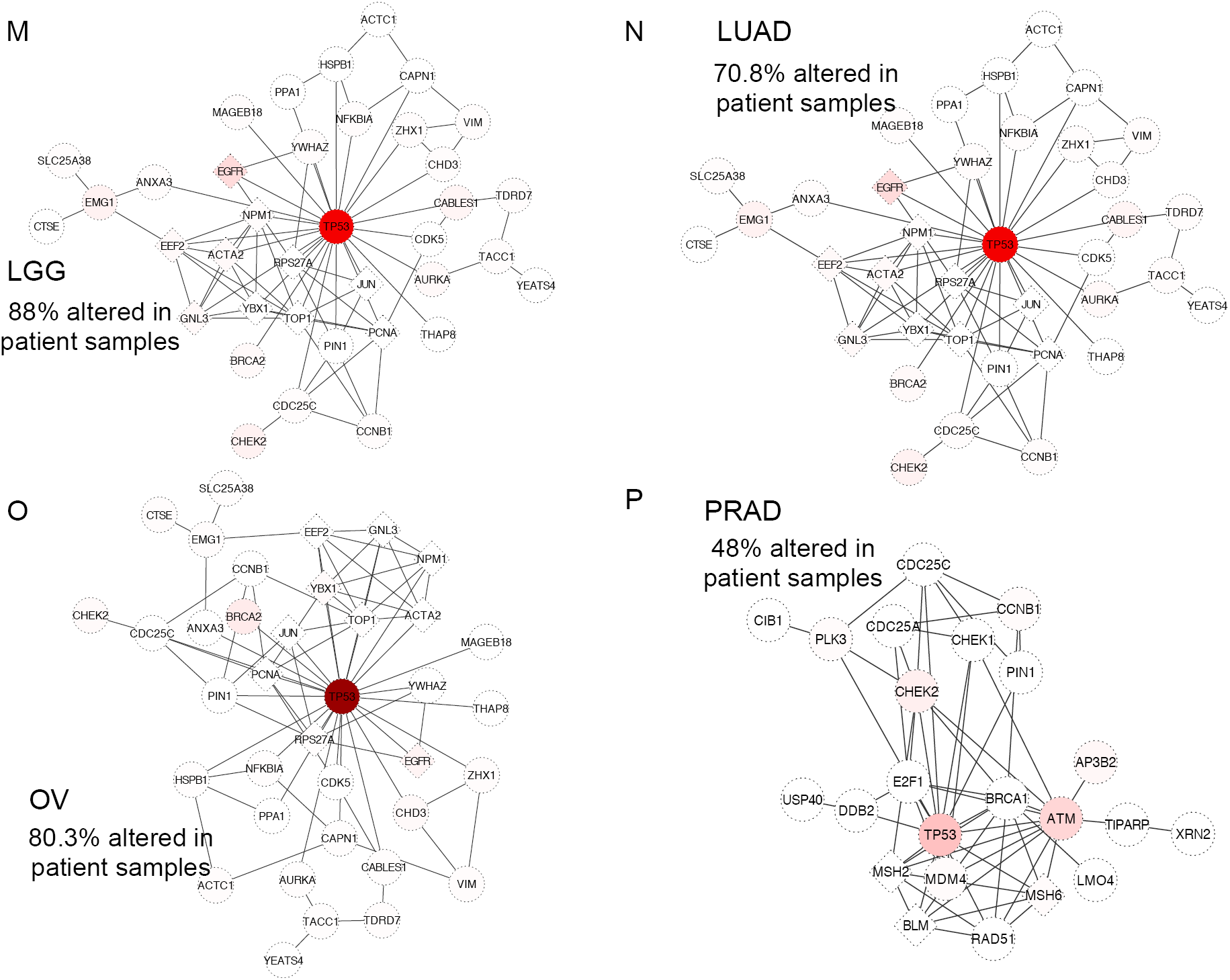
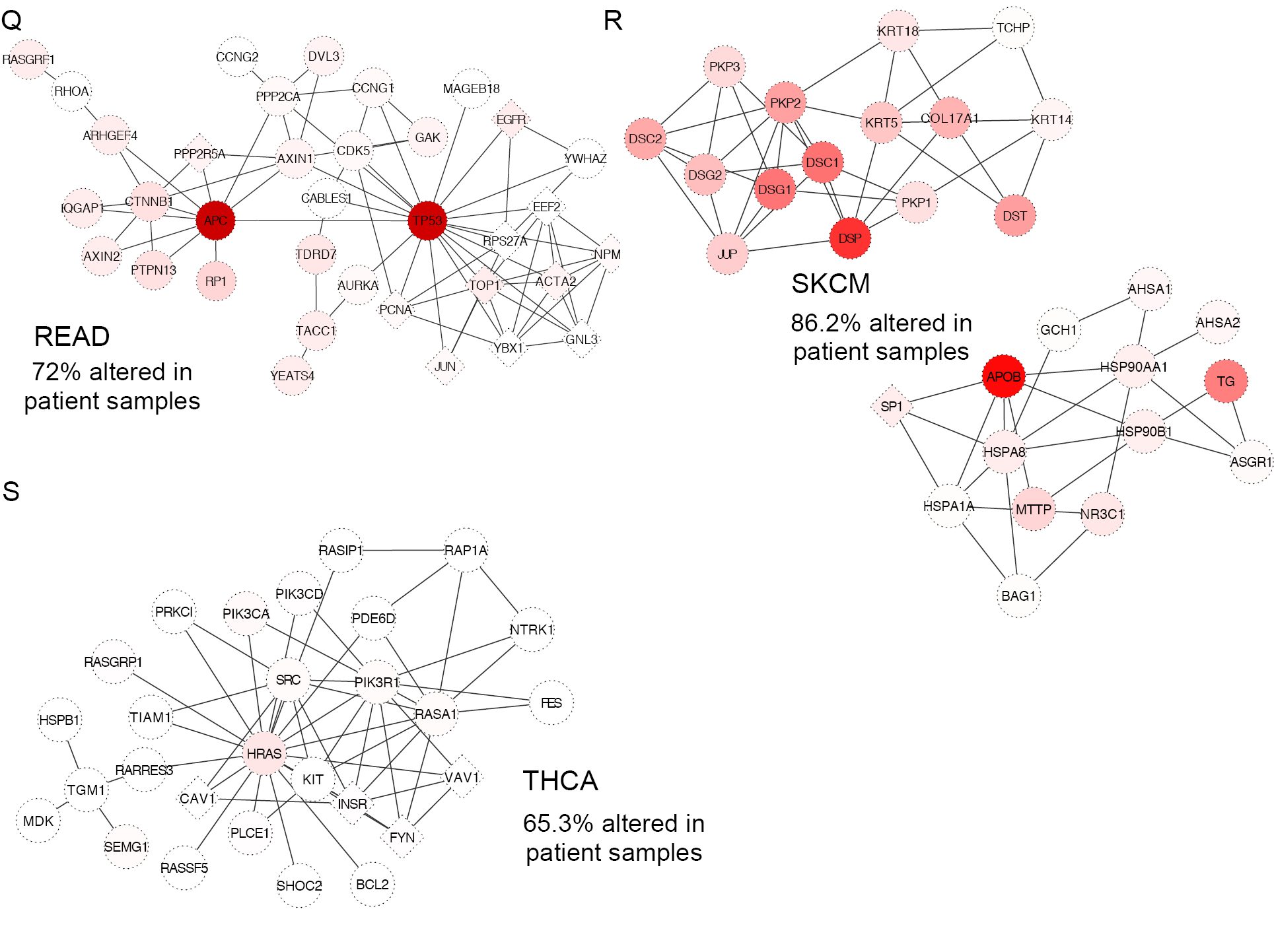
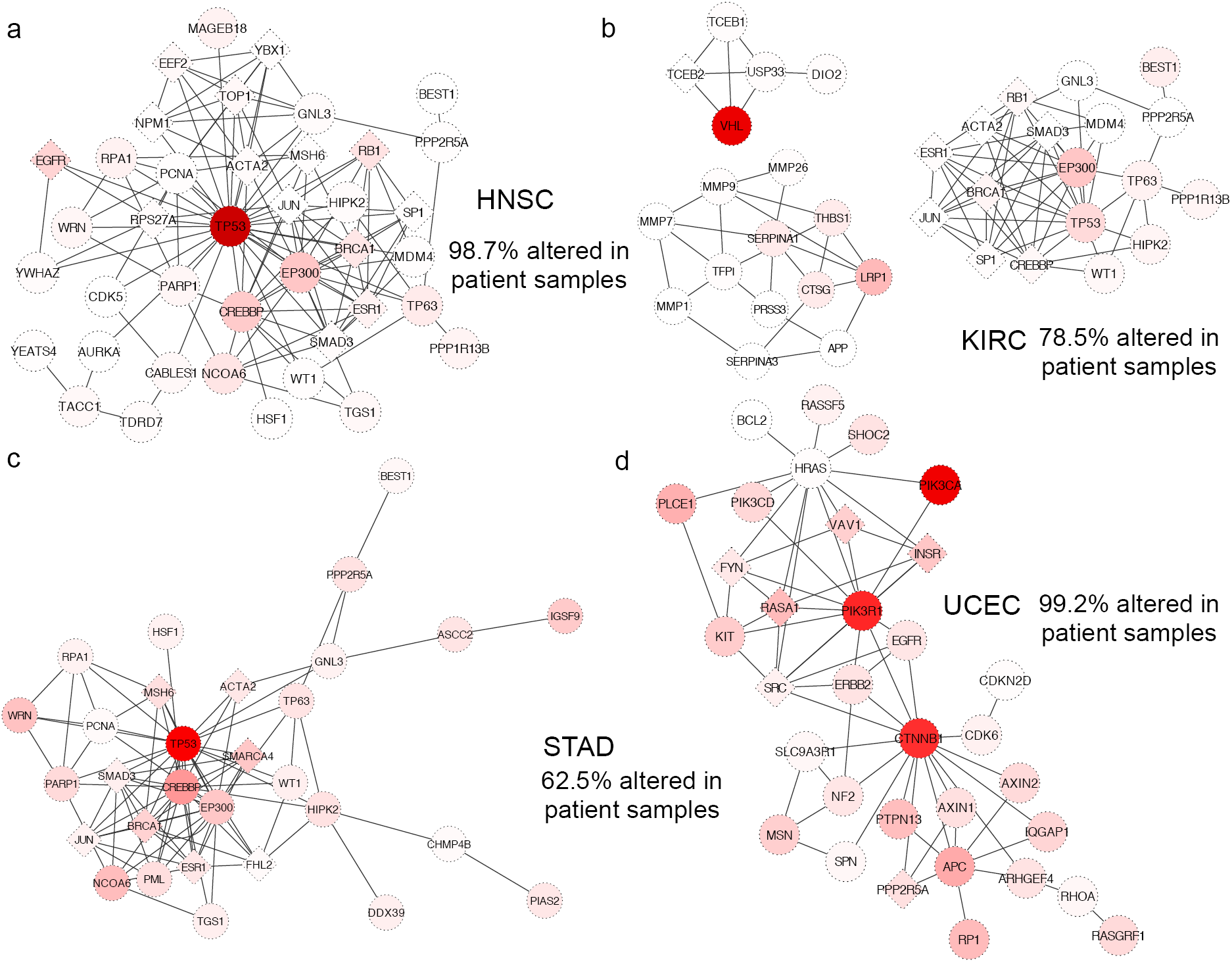
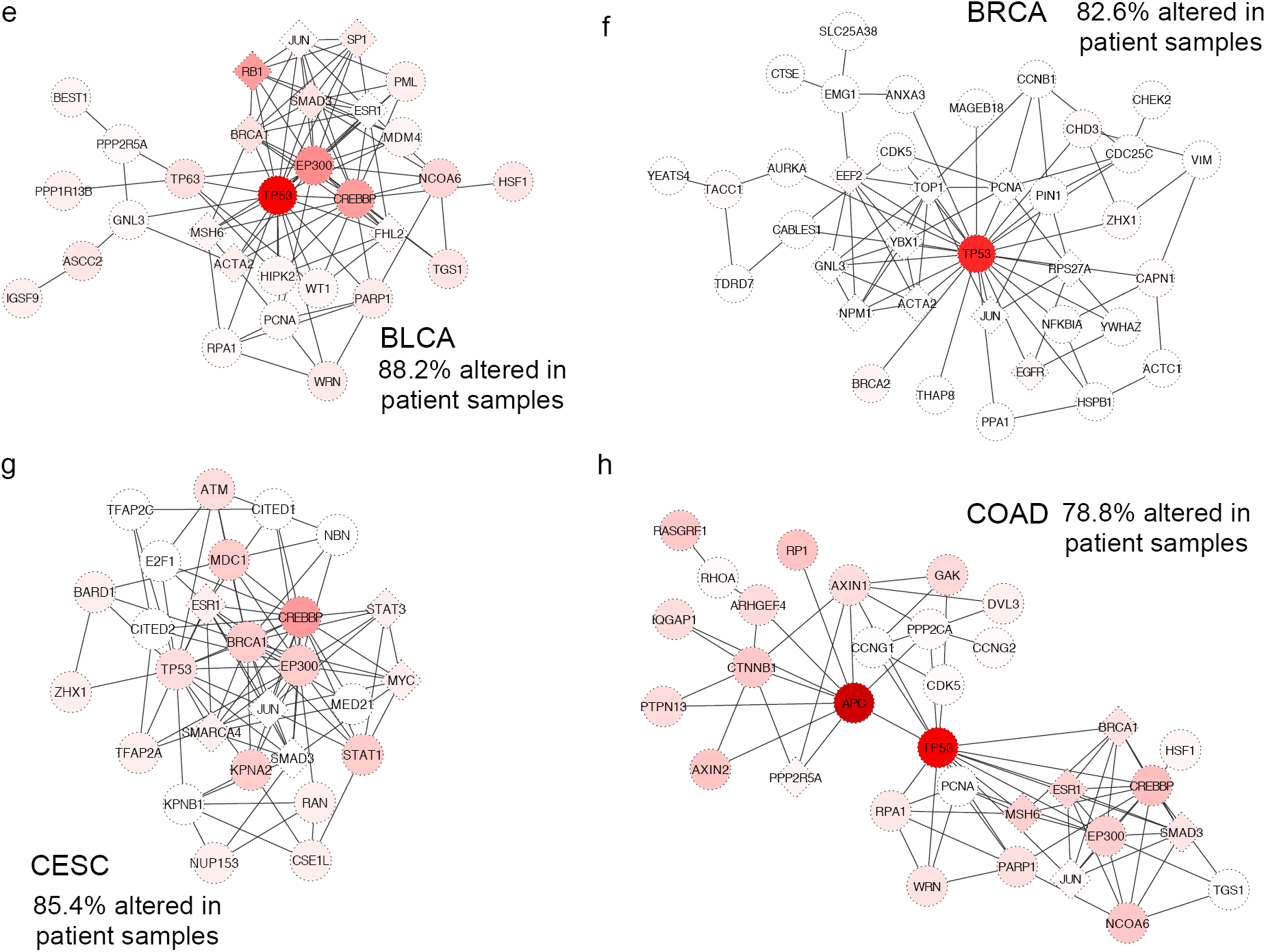
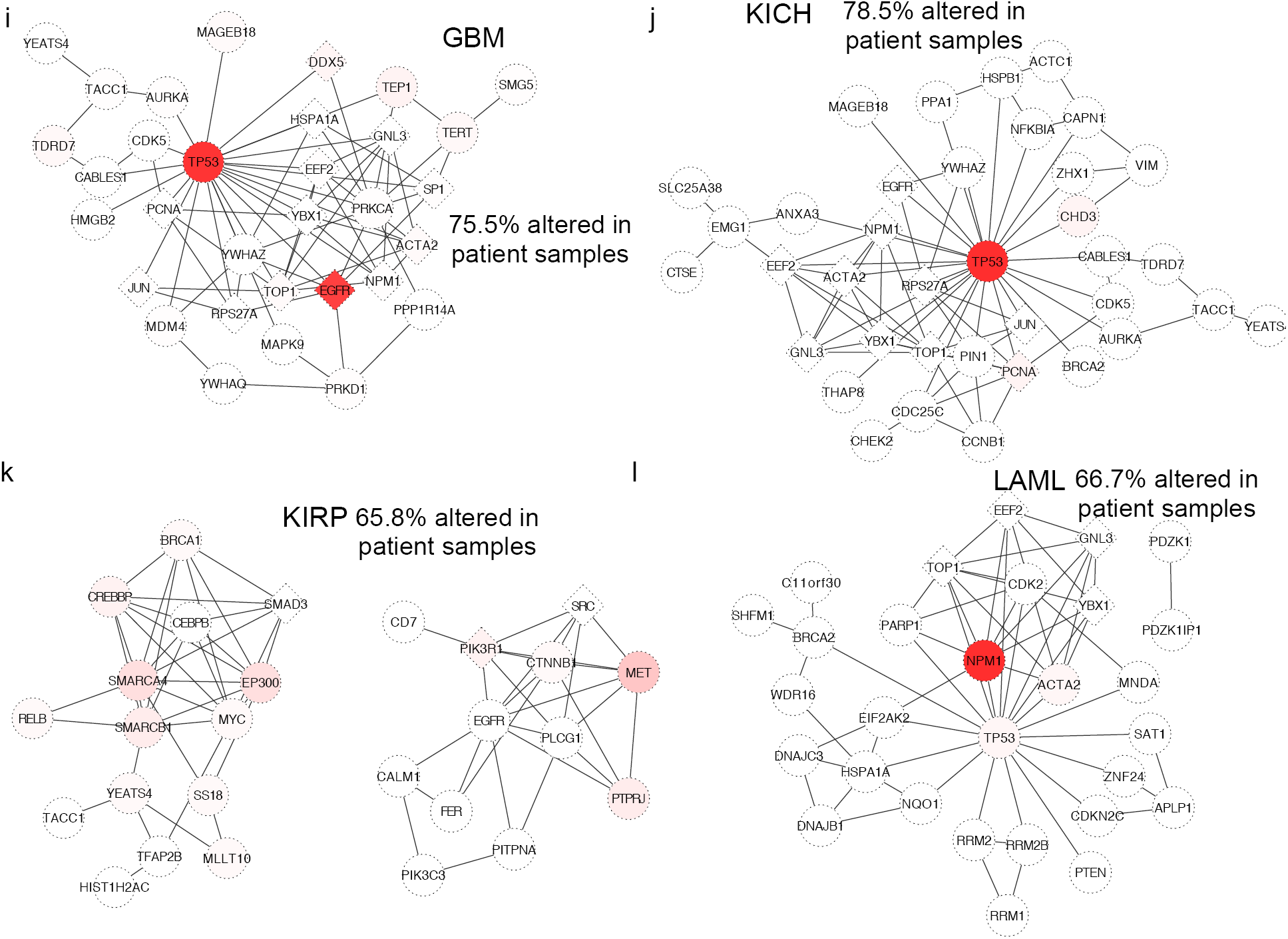
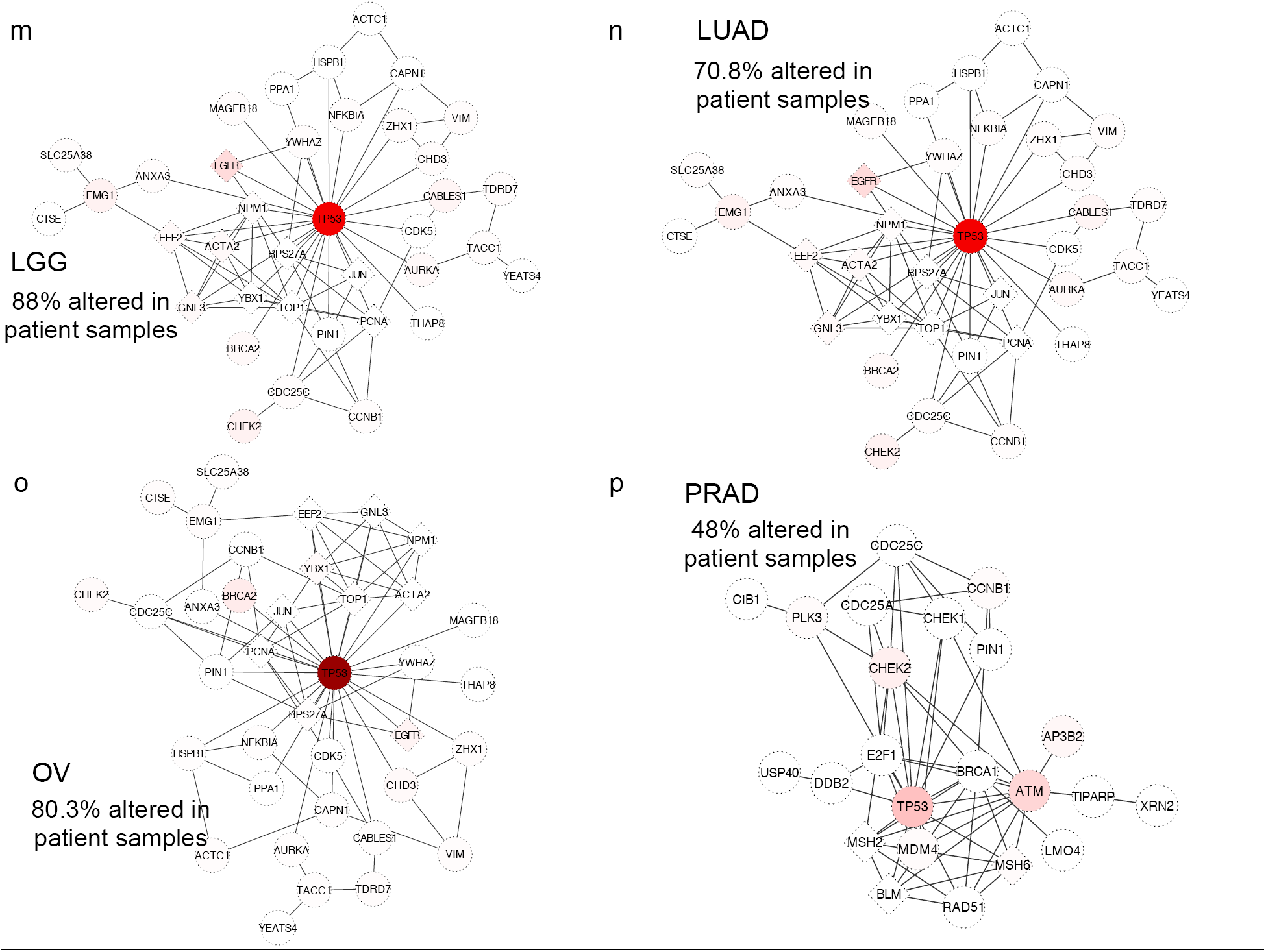
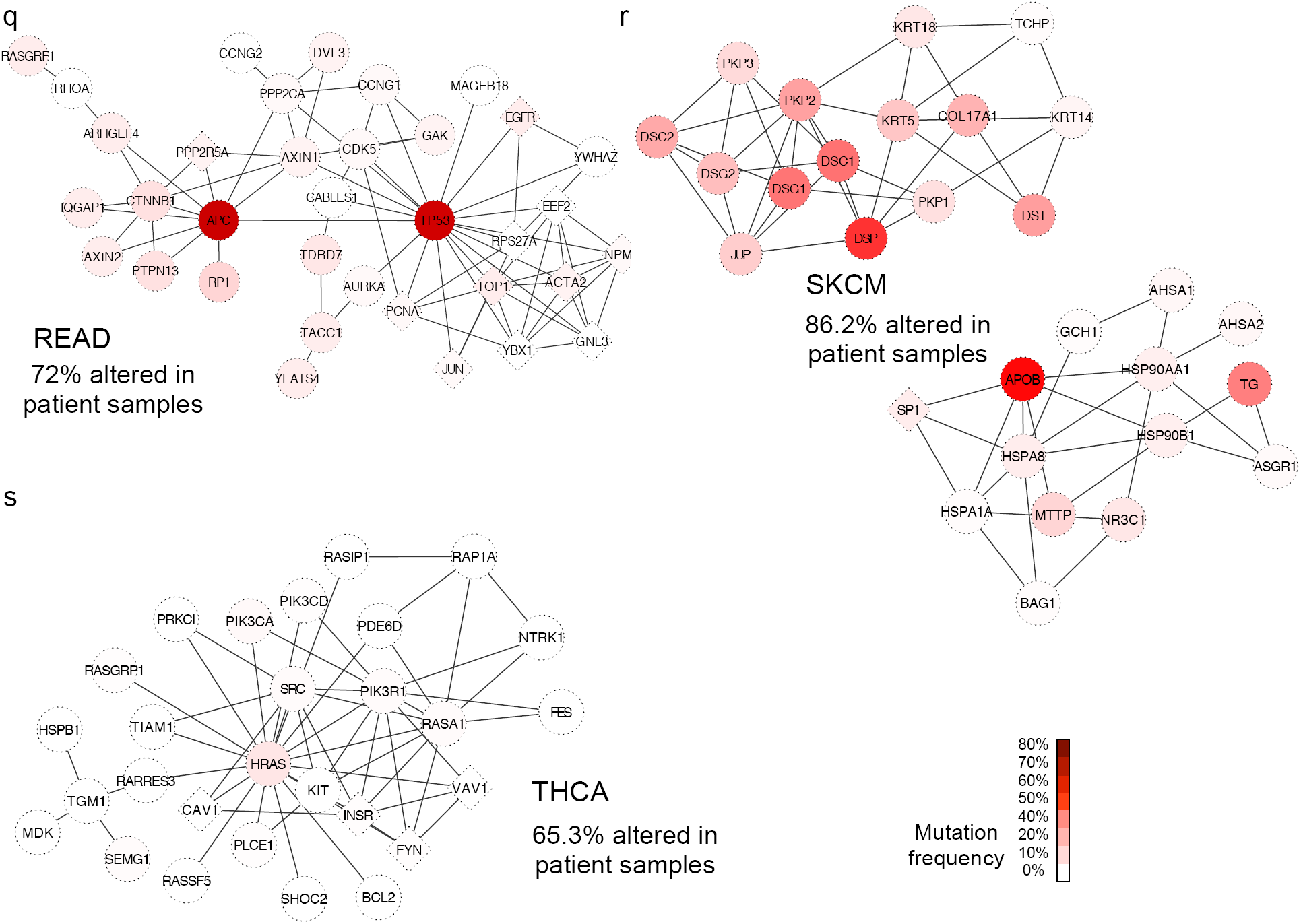
Supplementary Figure 1.

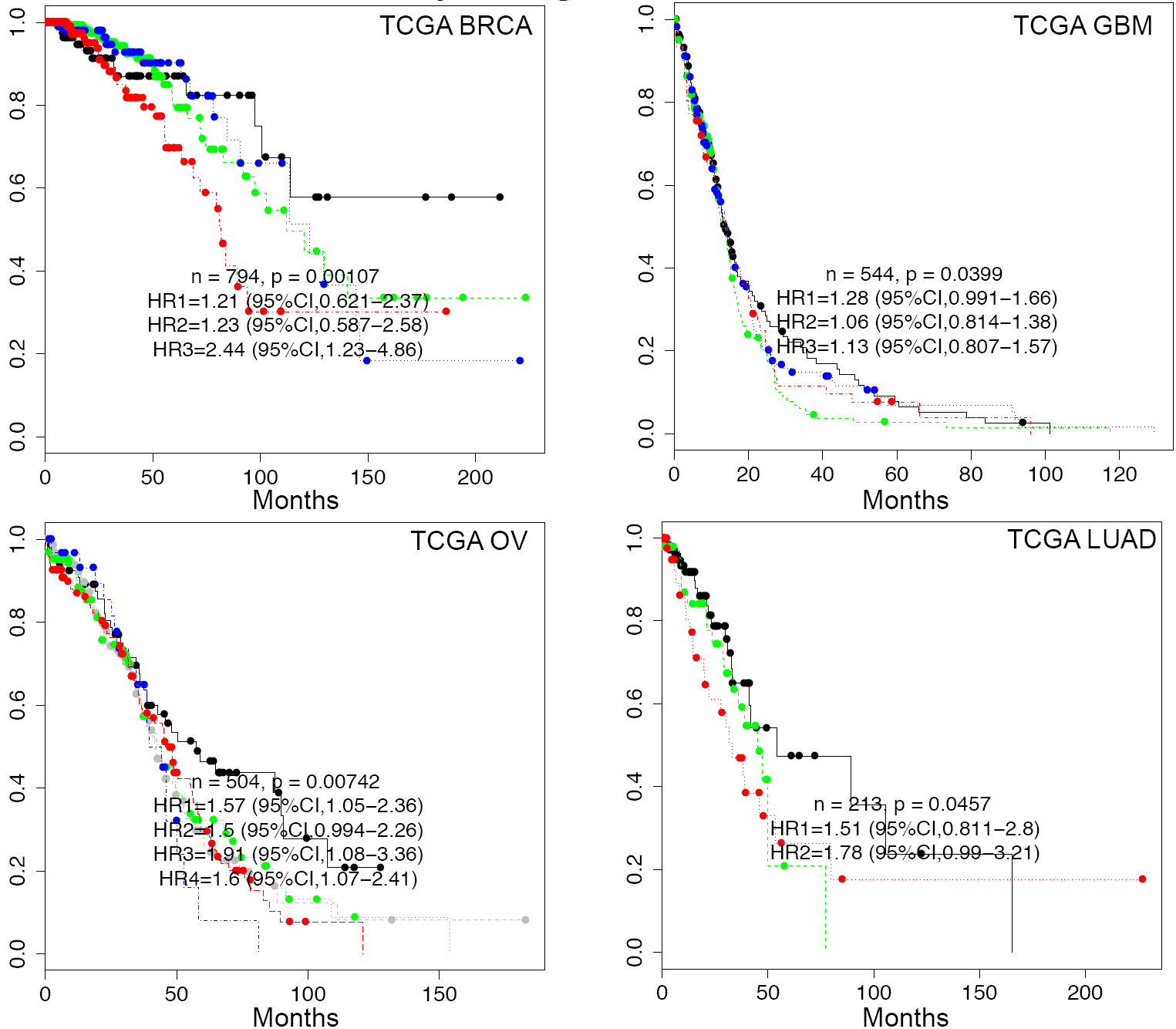
Supplementary Figure 2.

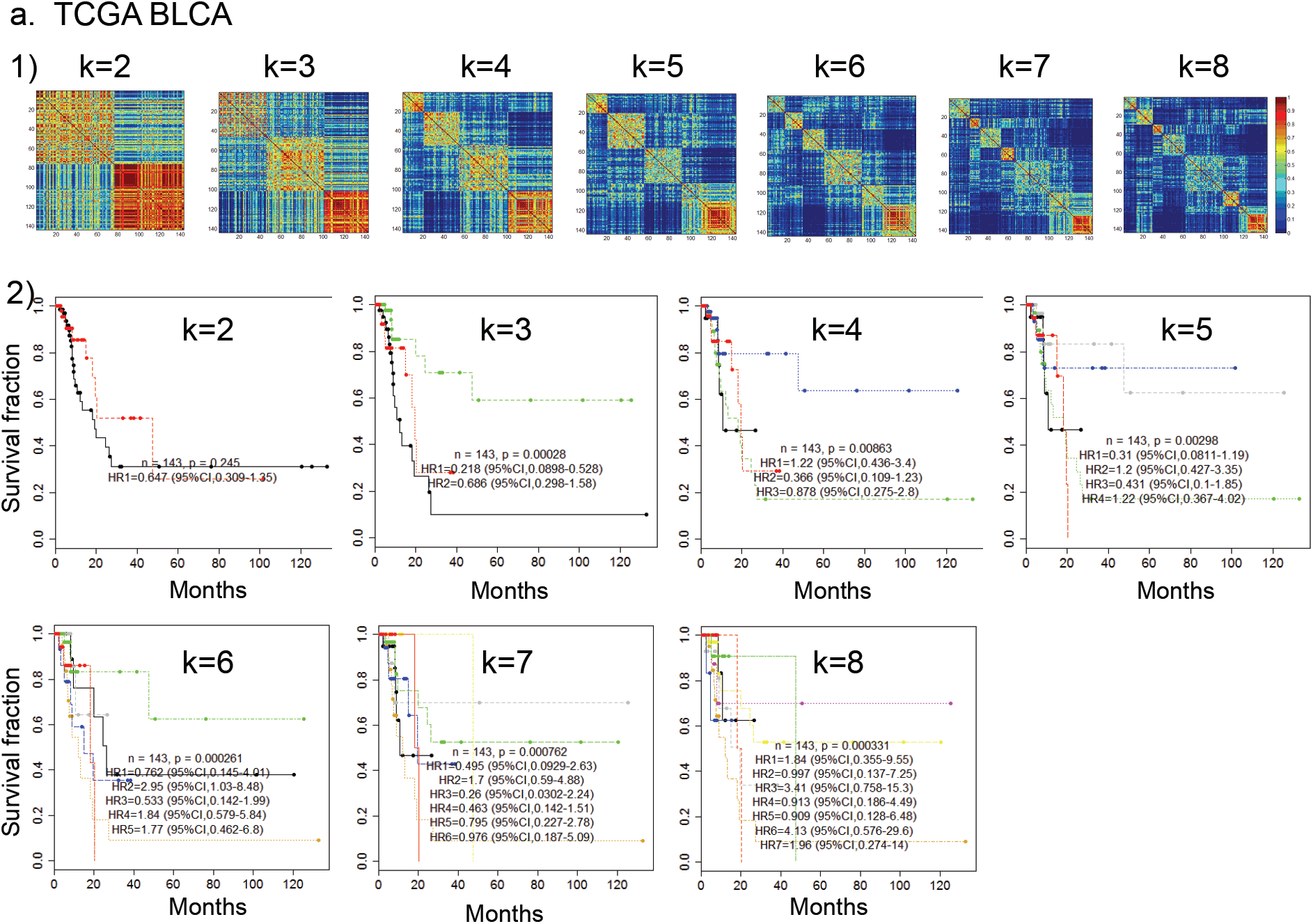
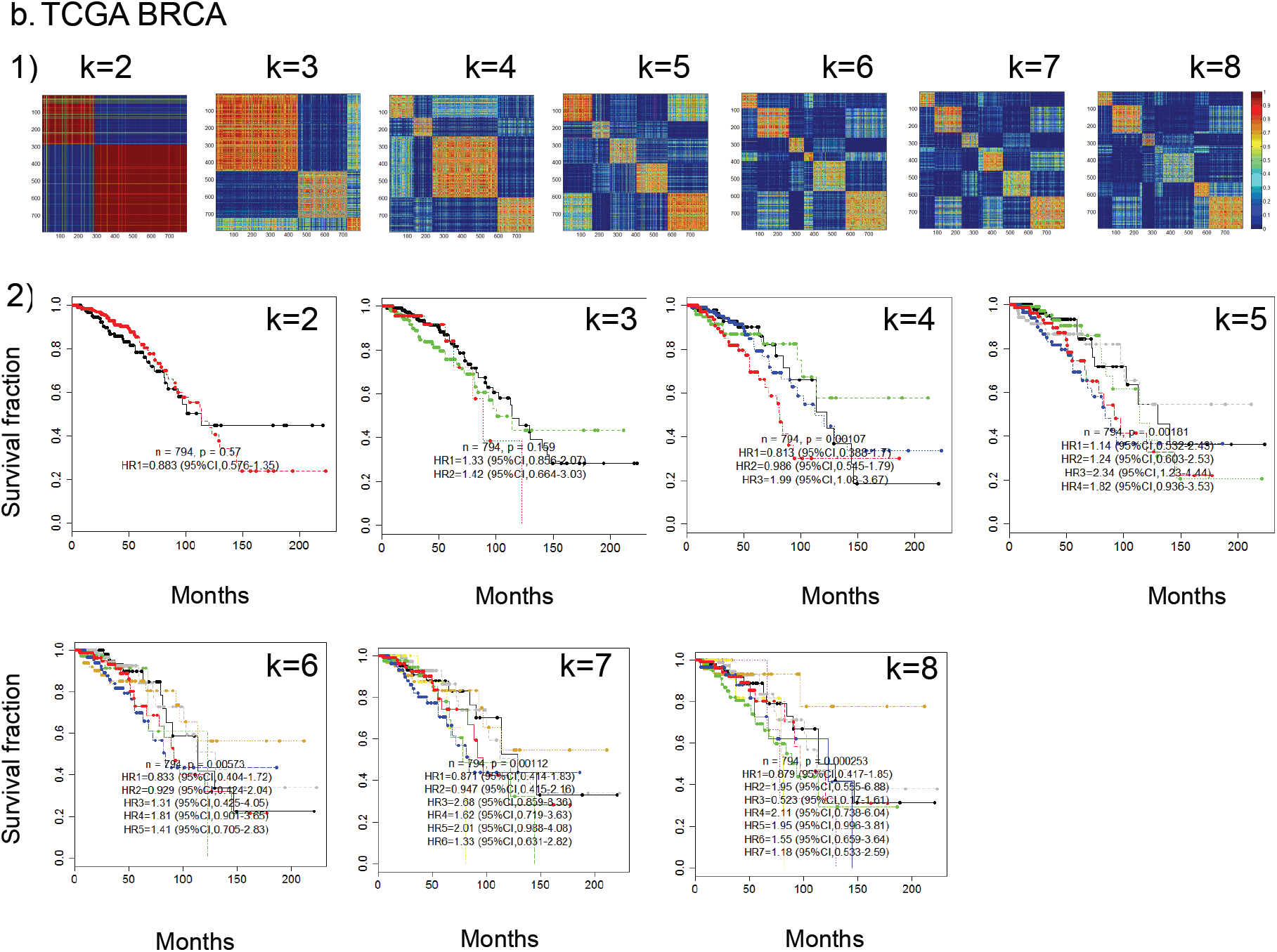
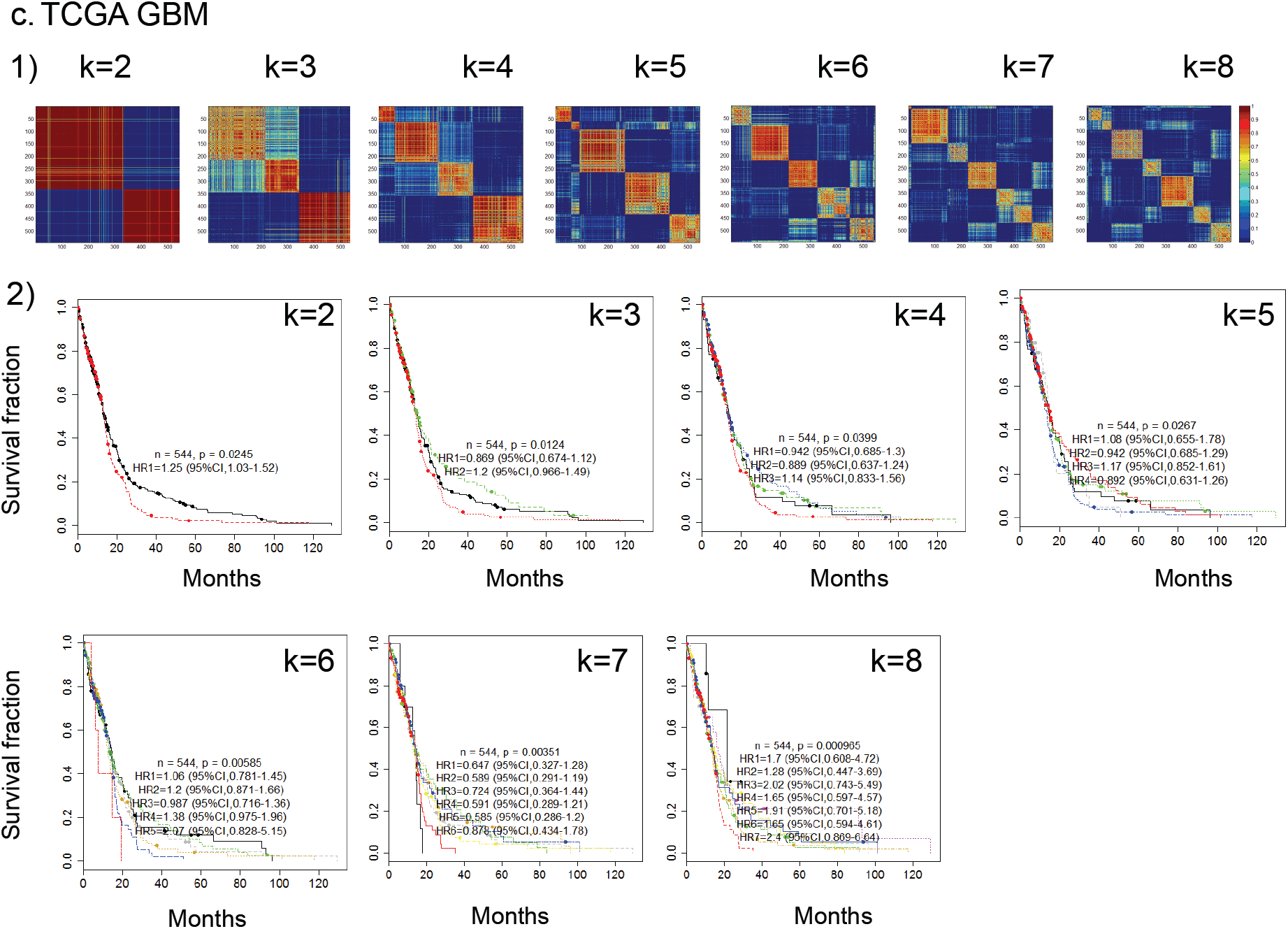
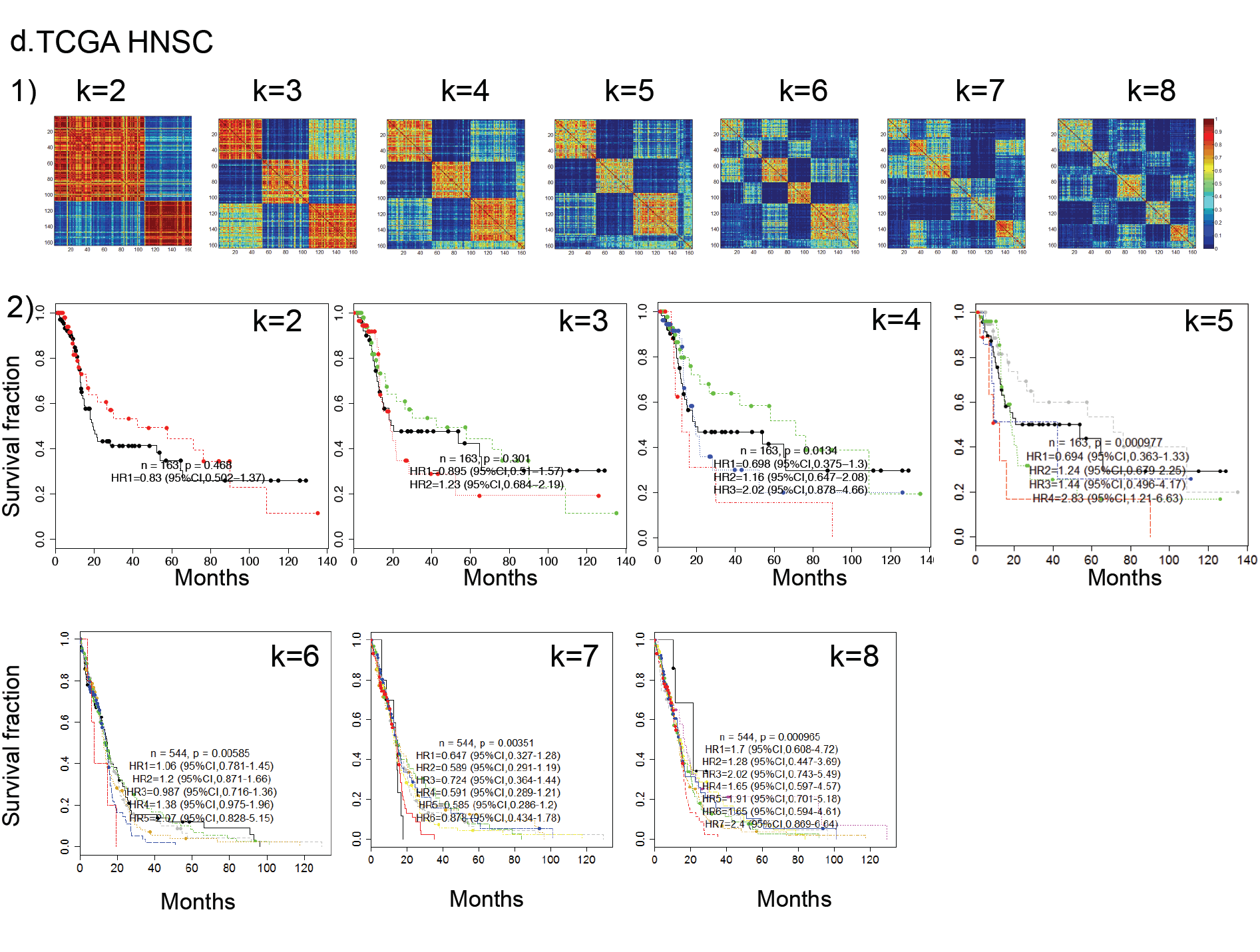
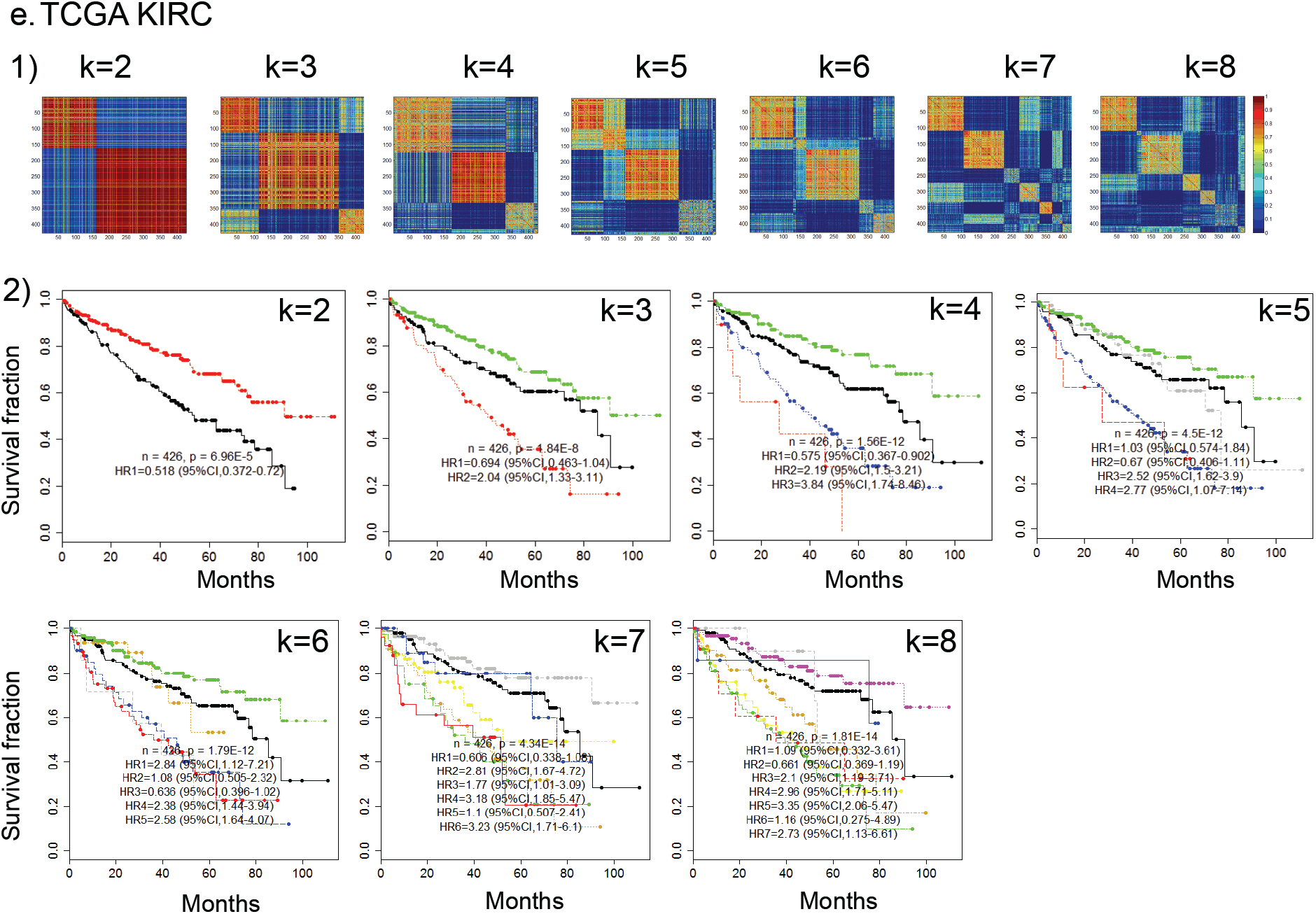
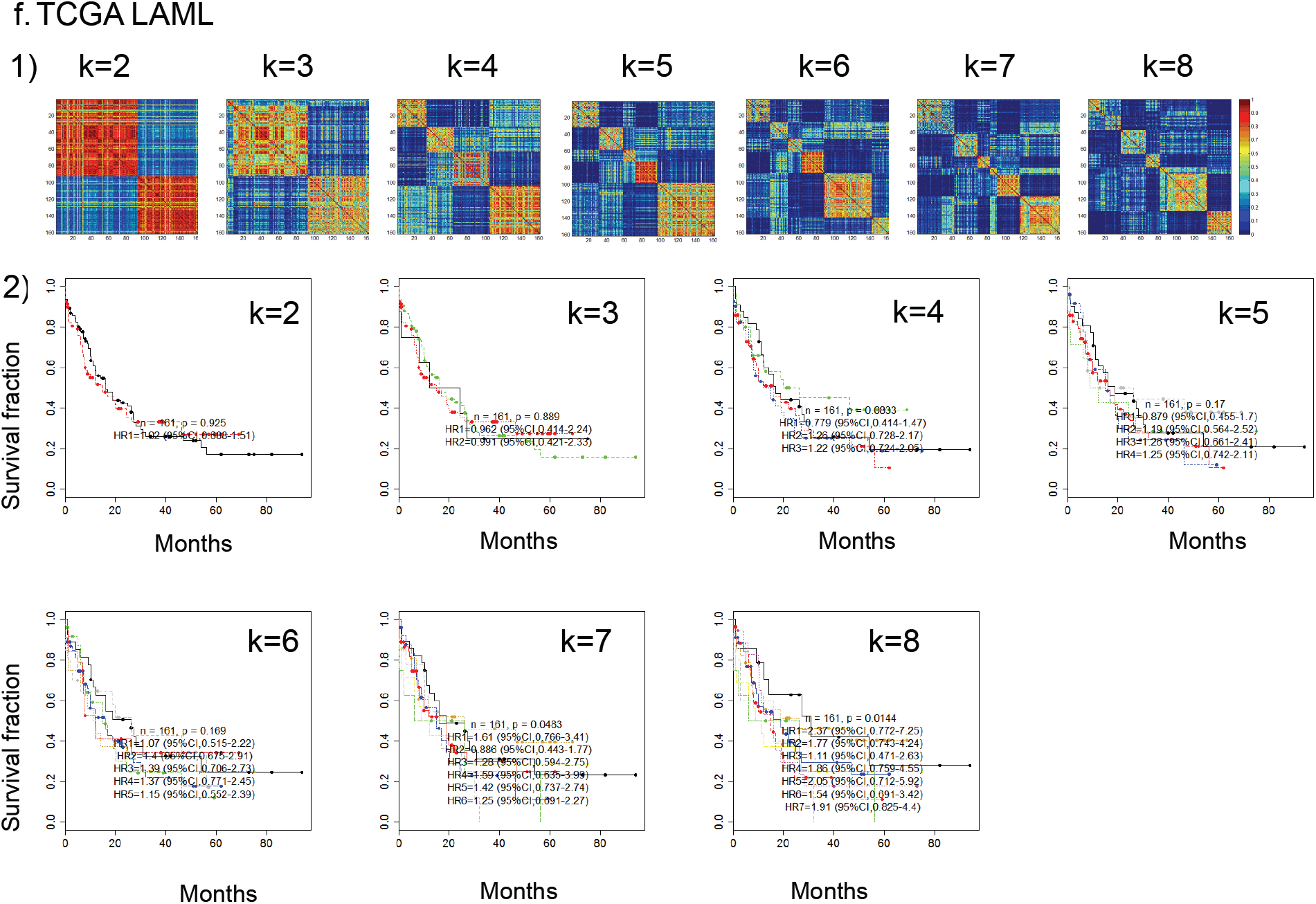
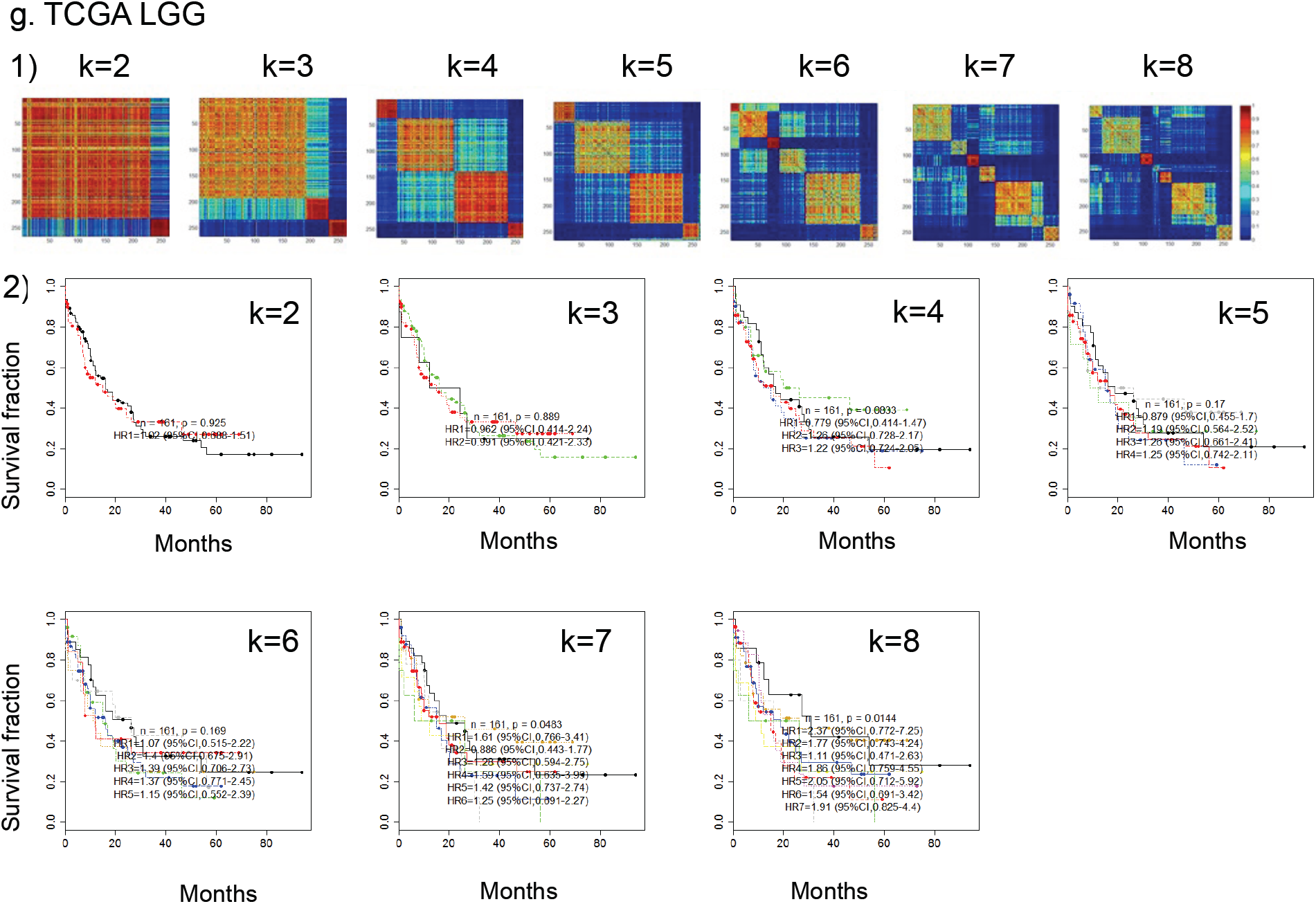
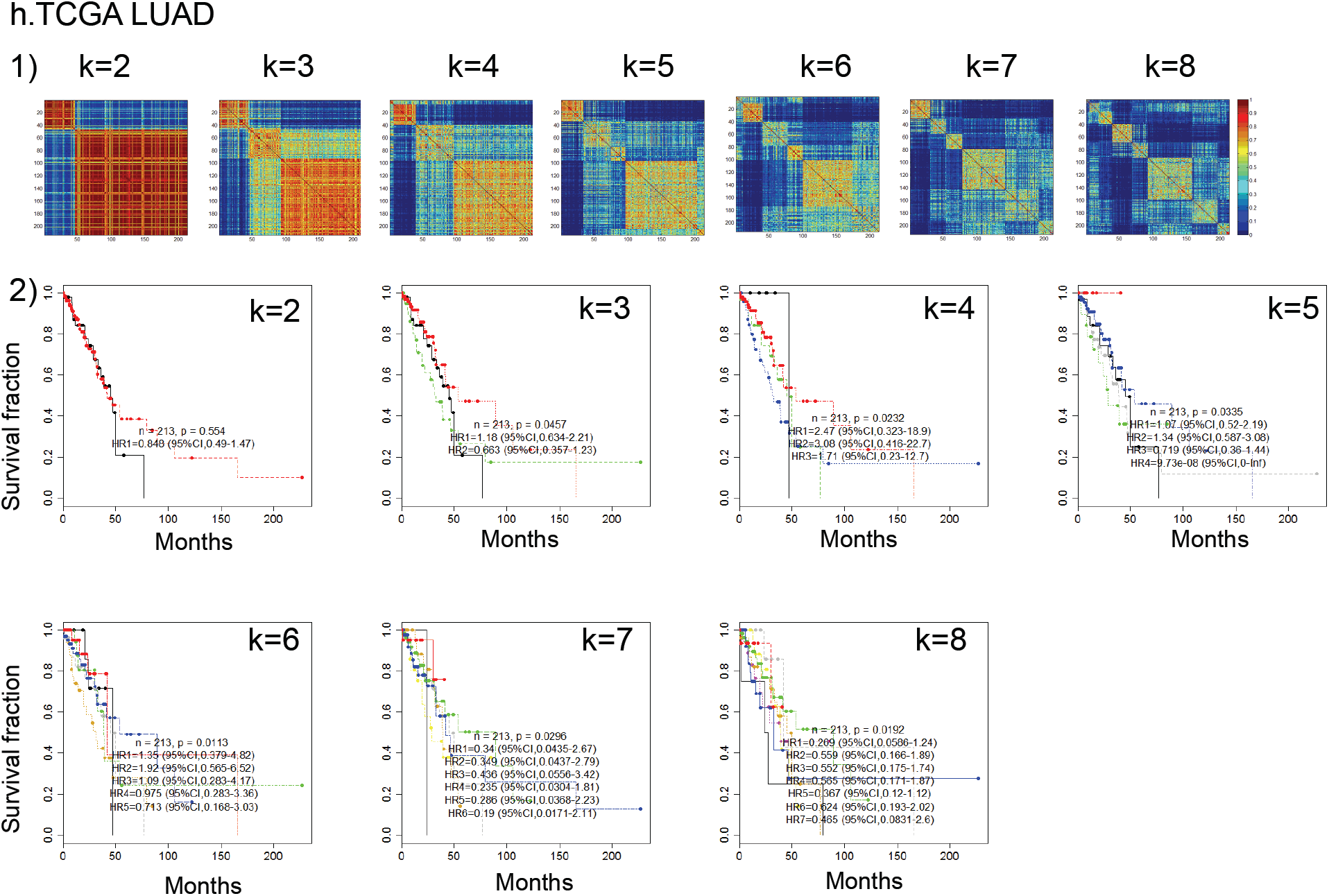
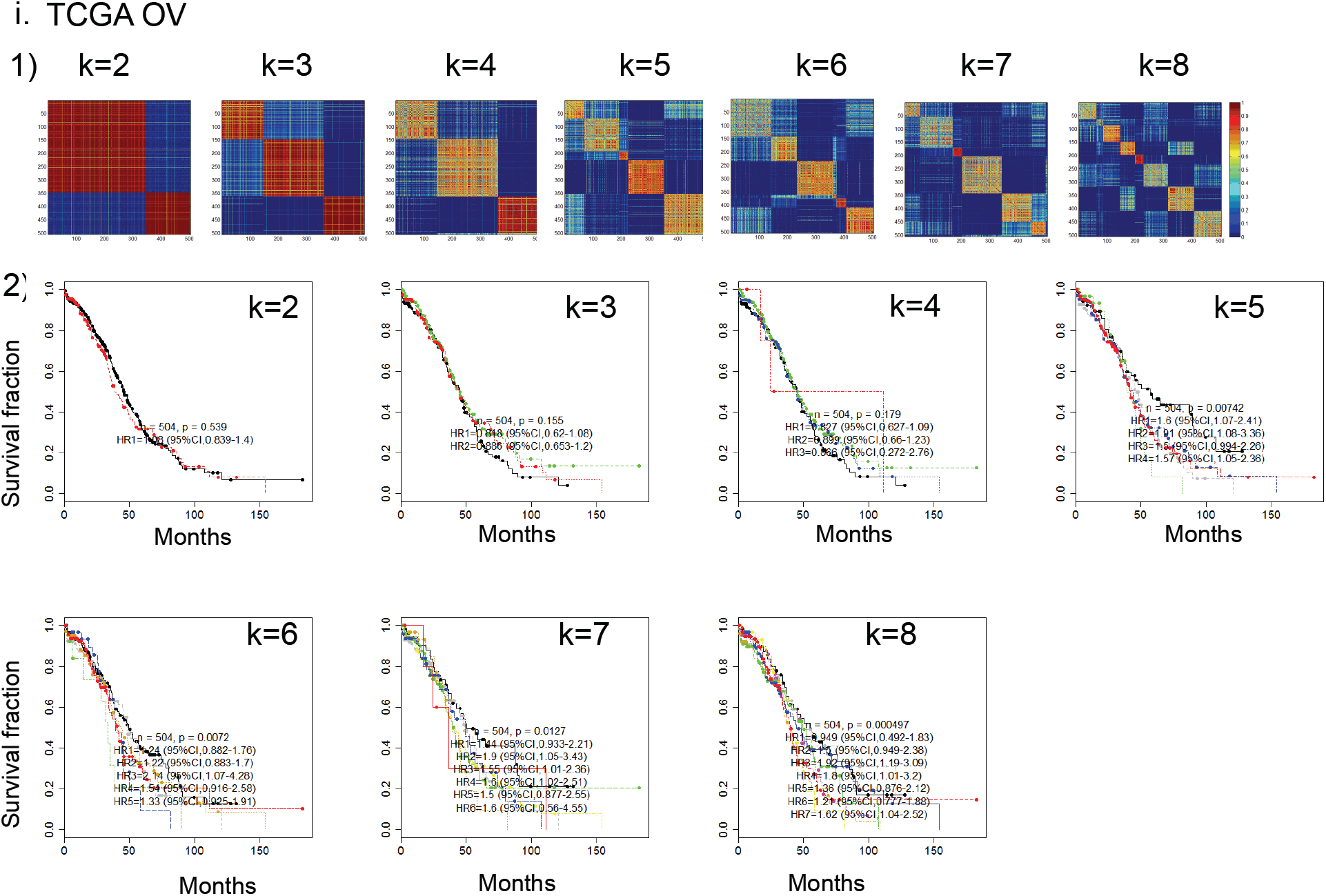
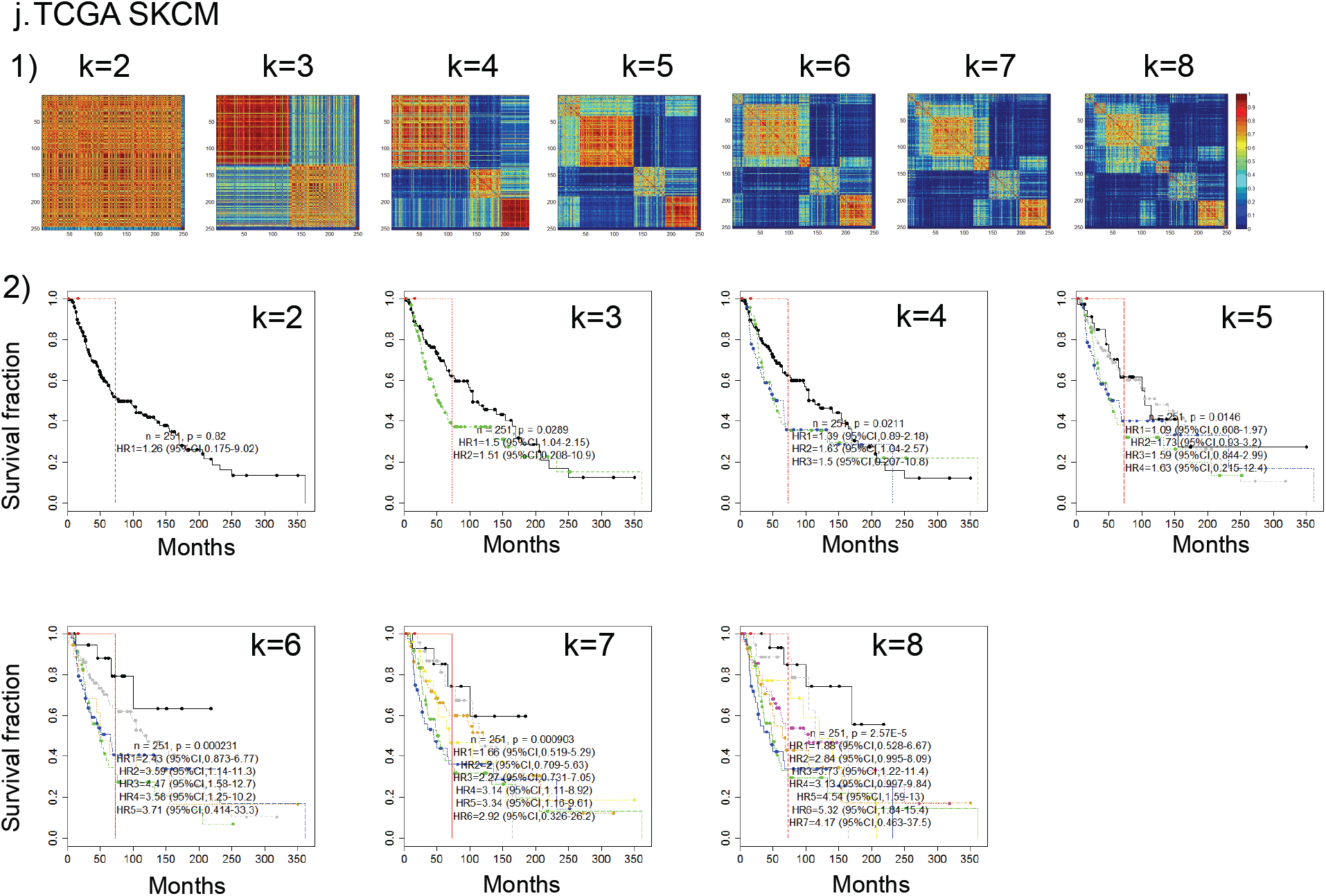
Supplementary Figure 3 KM plots.

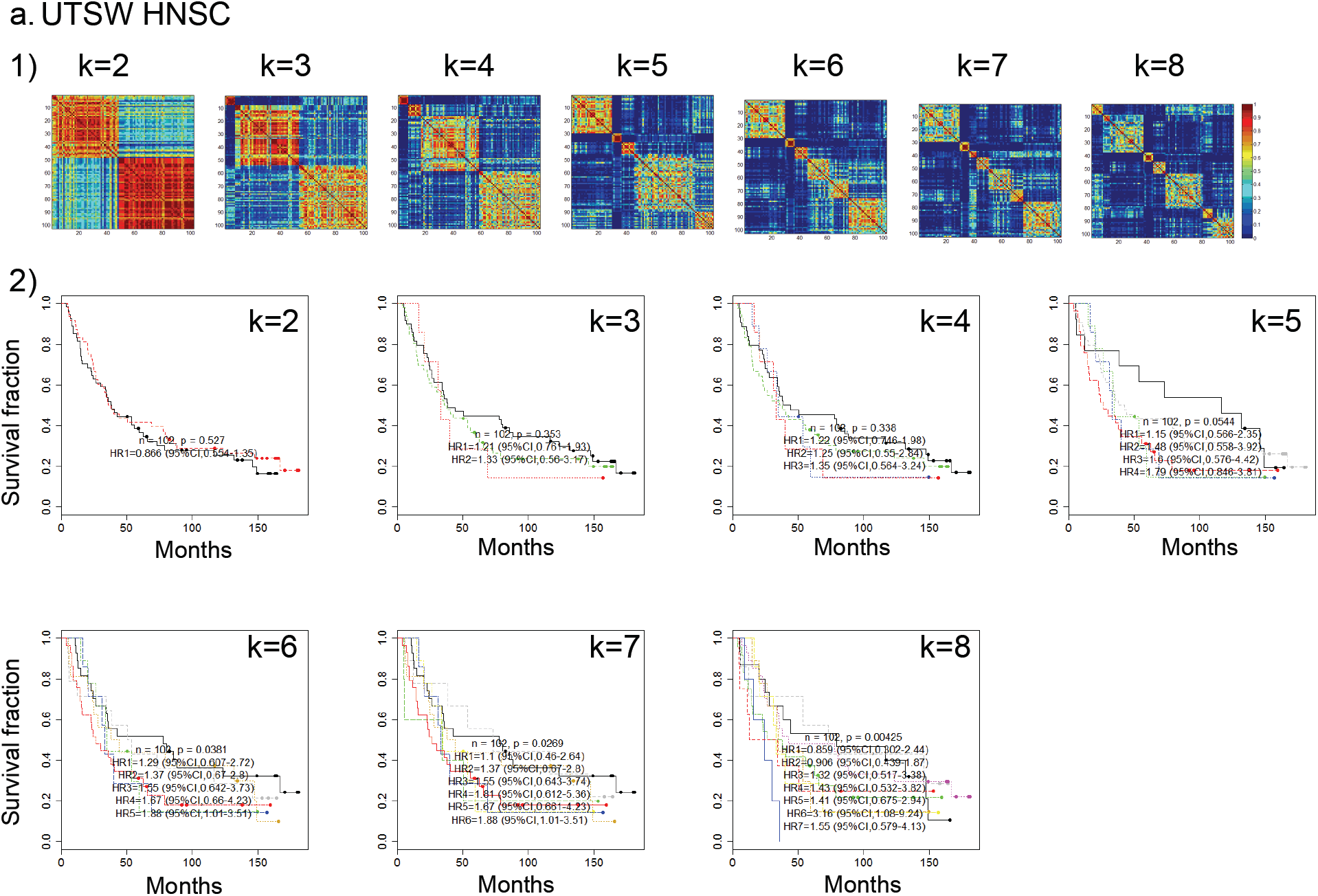
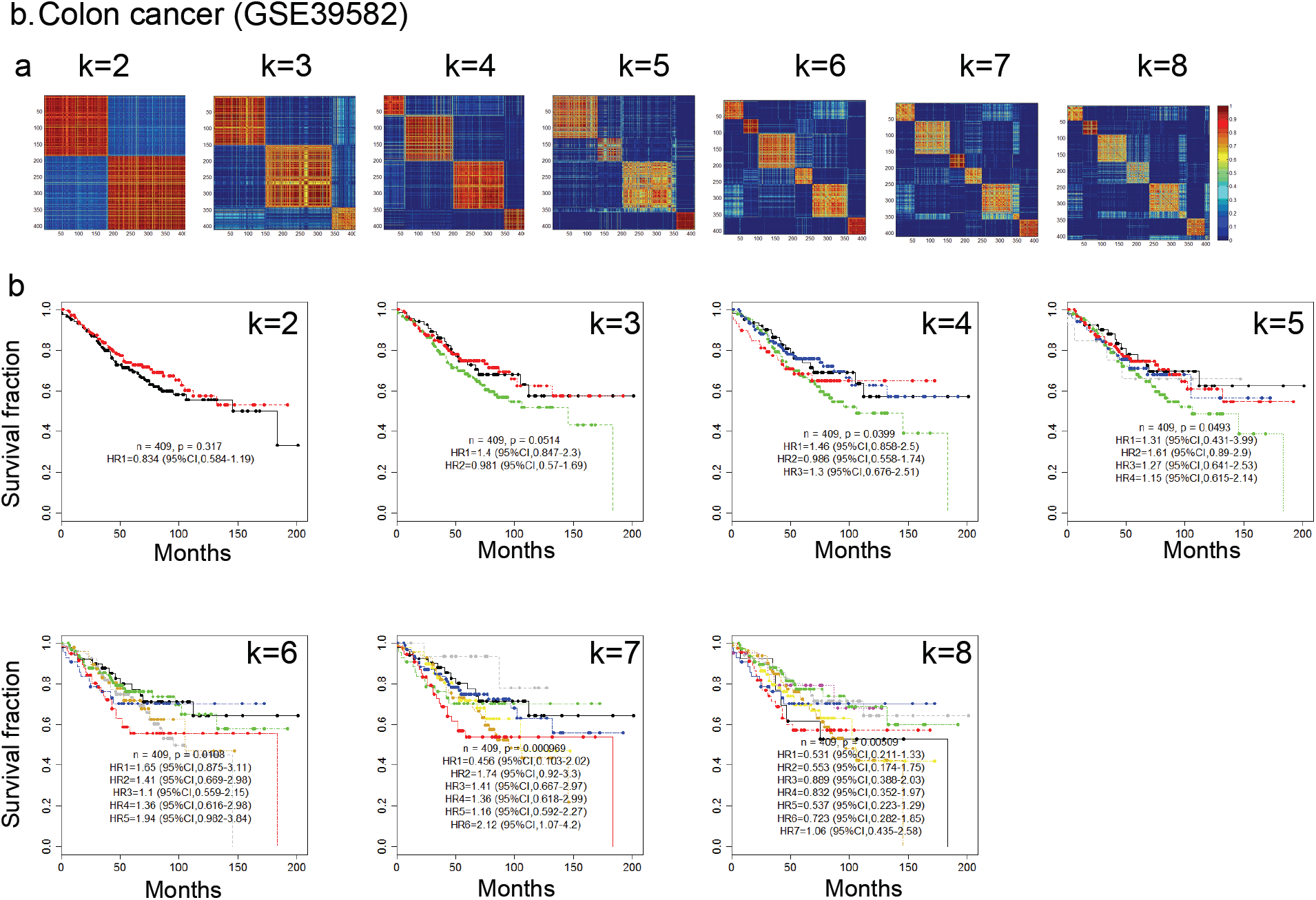
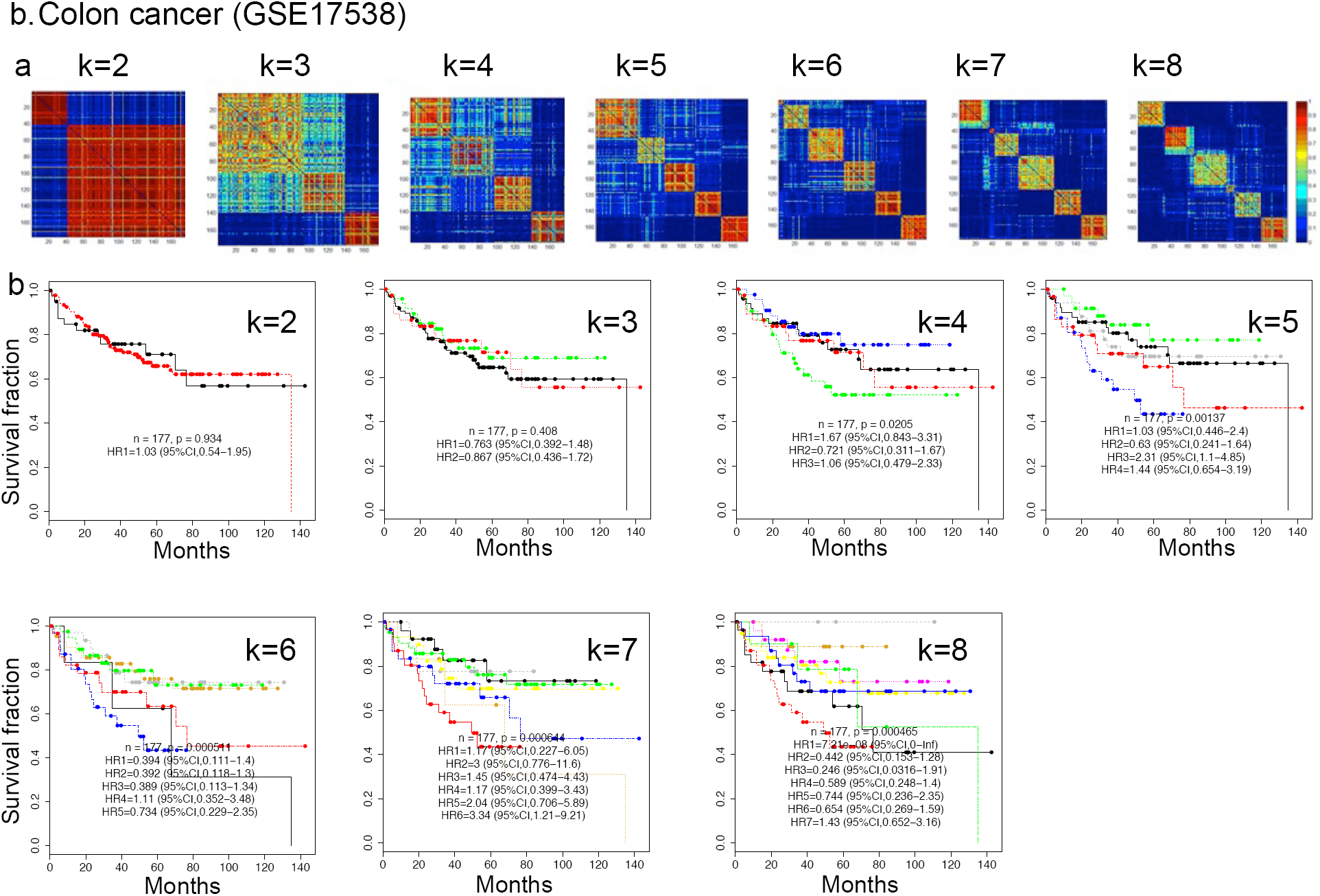
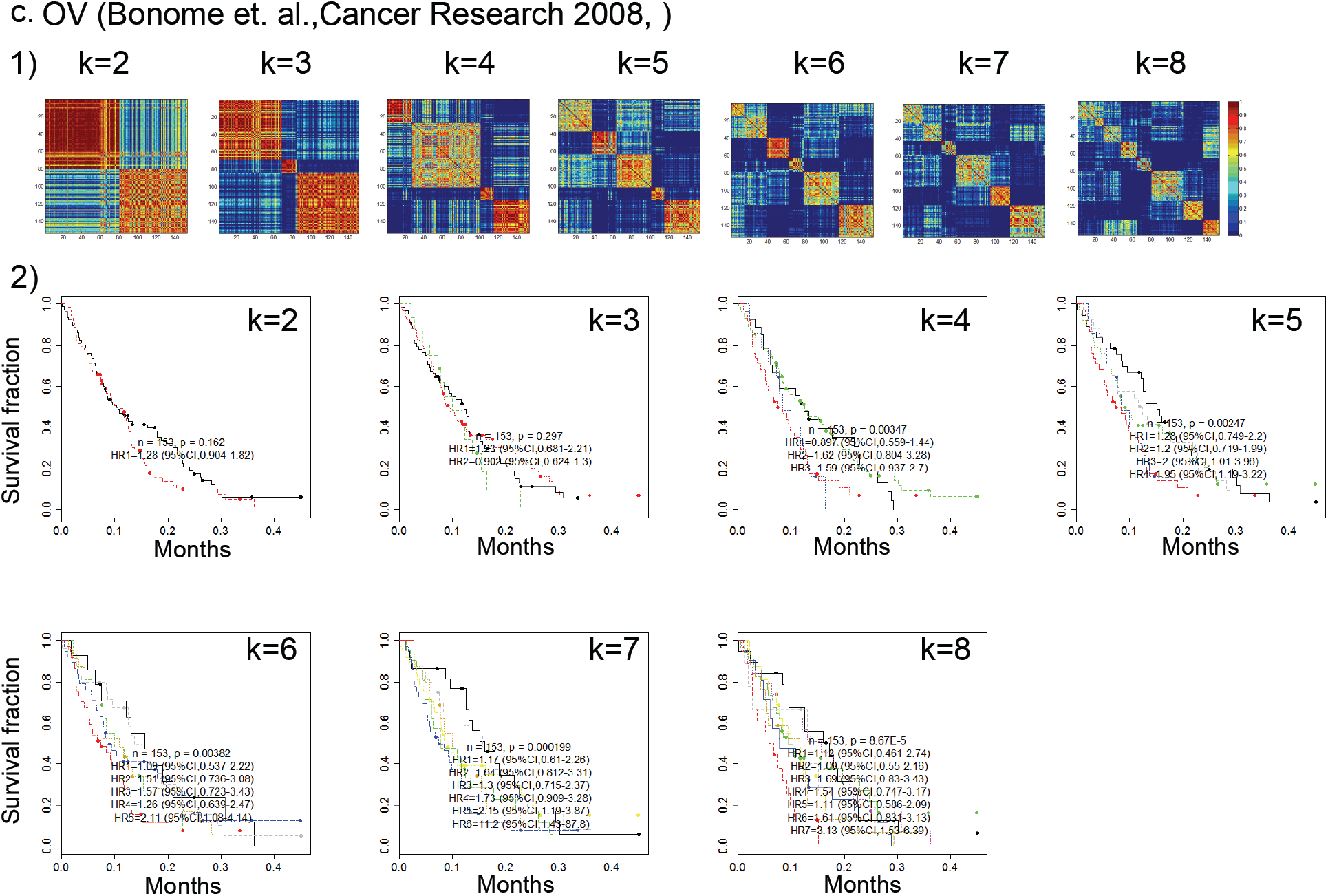
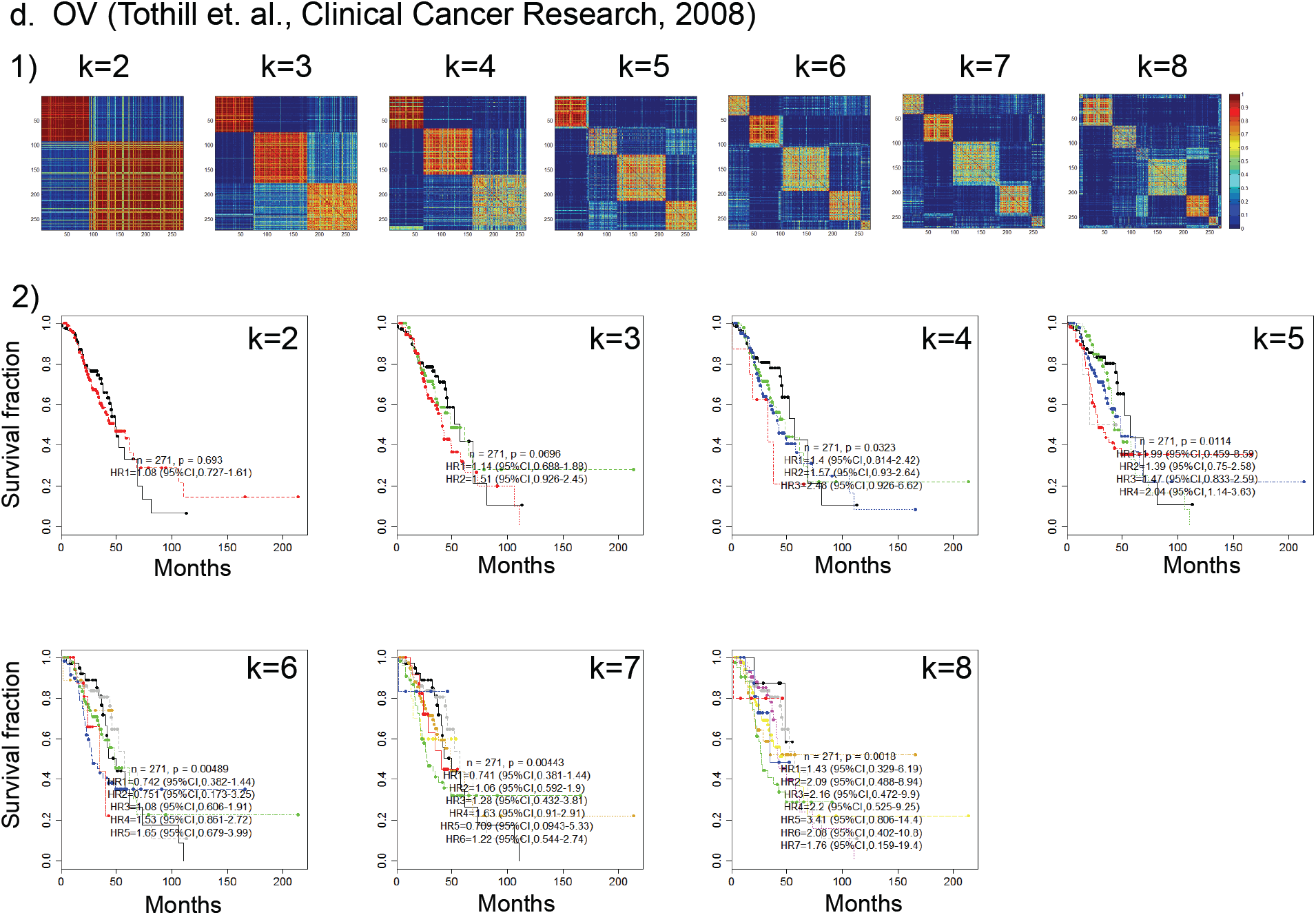
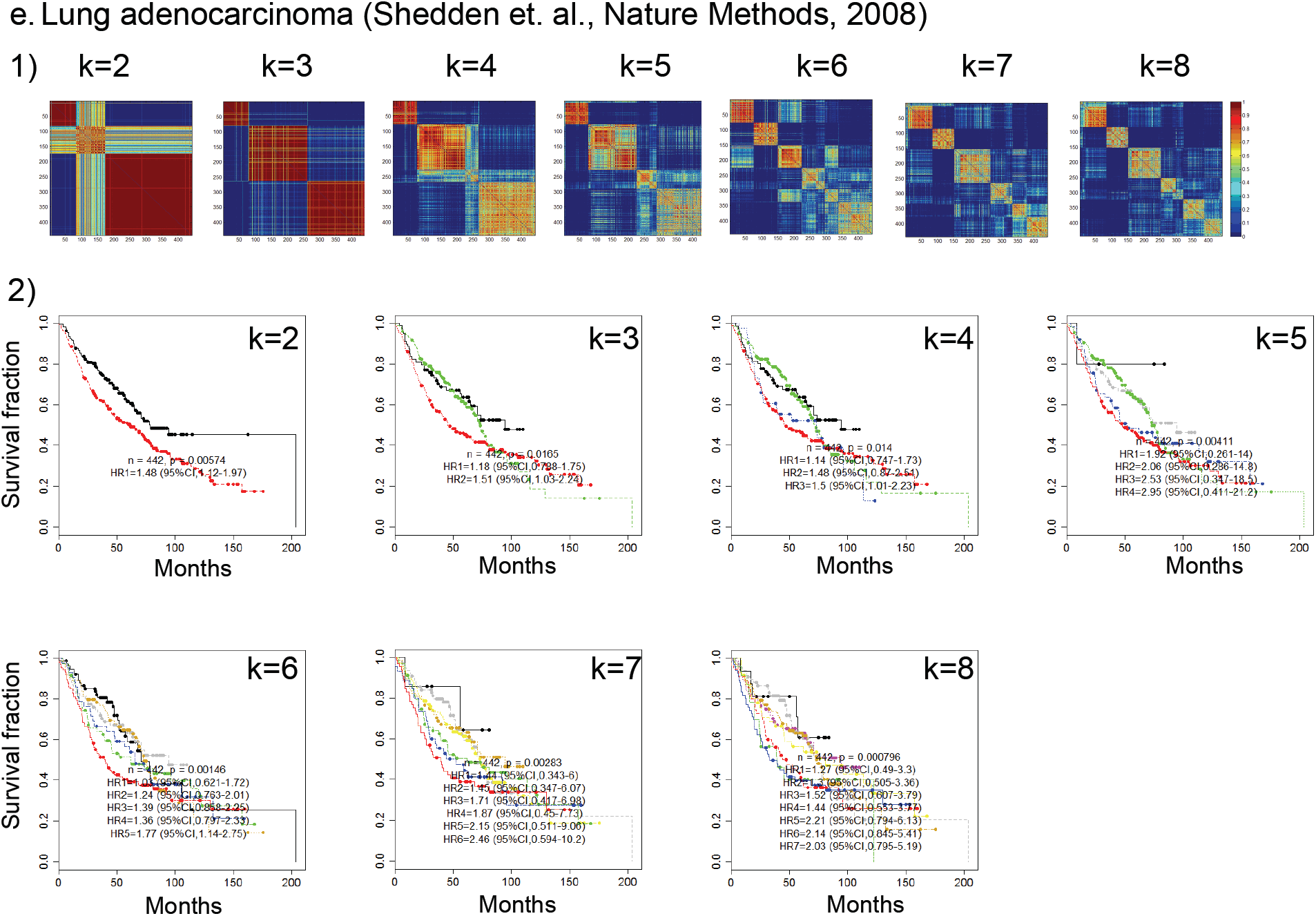
Supplementary Figure 4 Independent dataset.

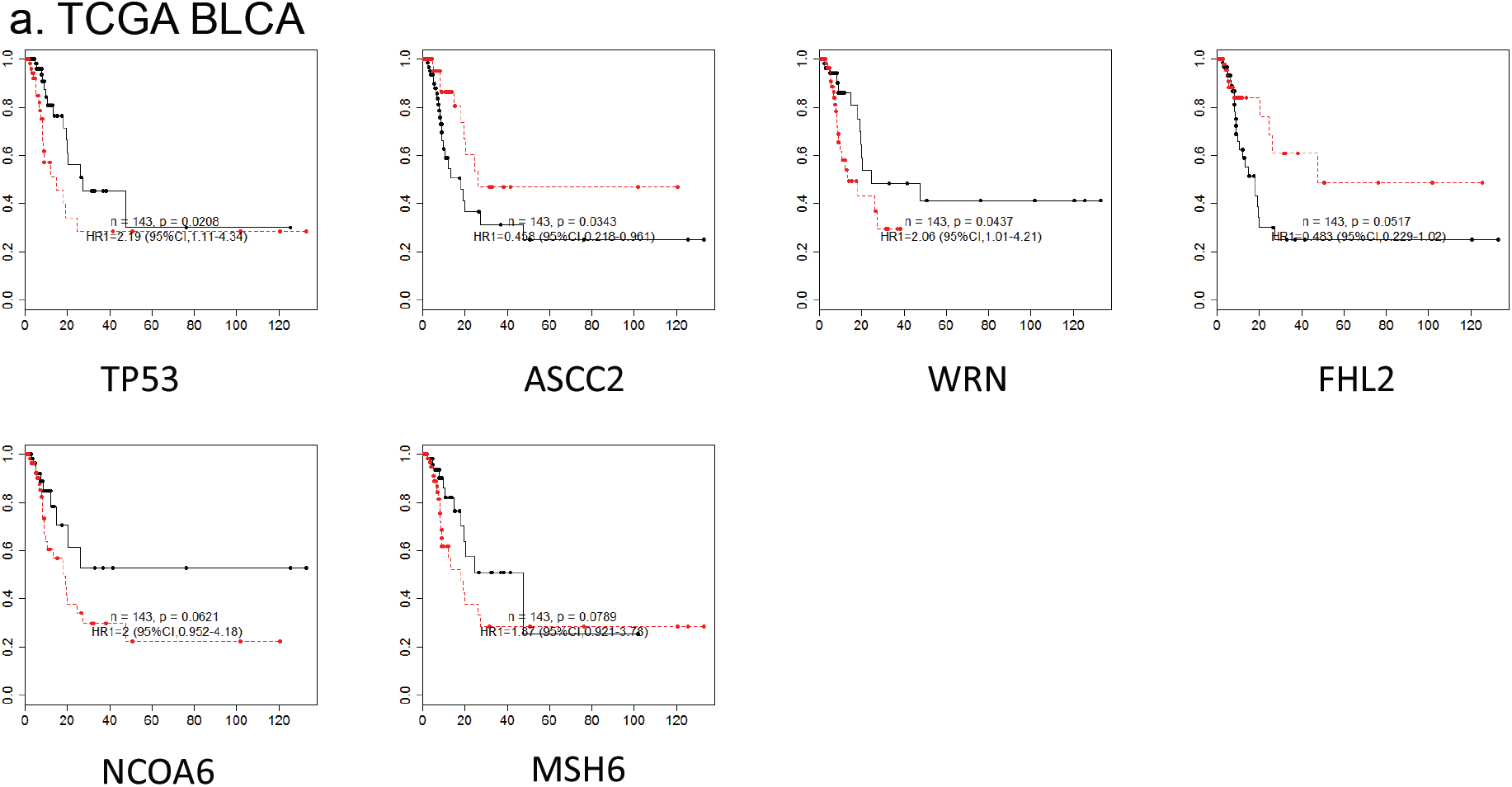
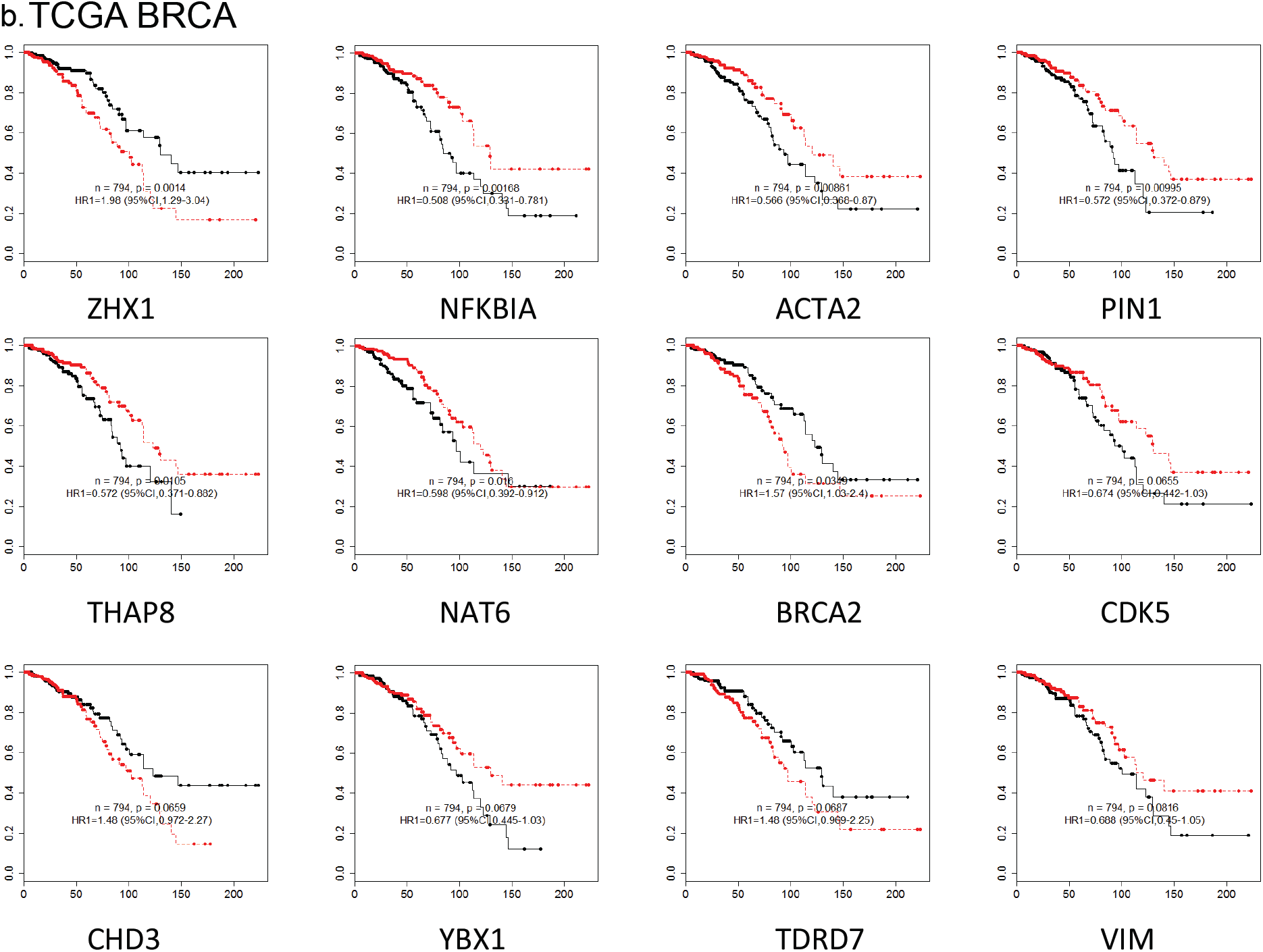
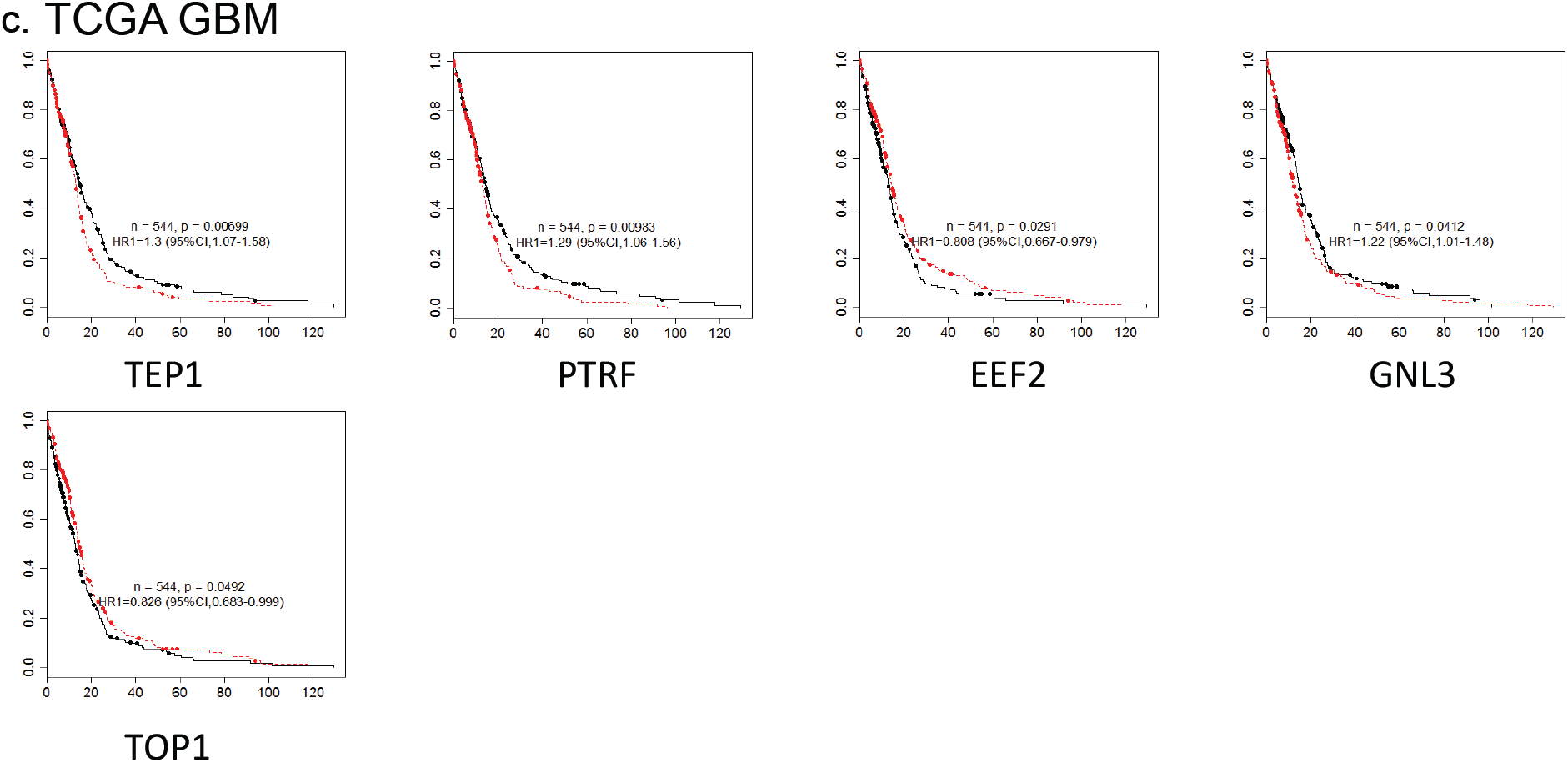
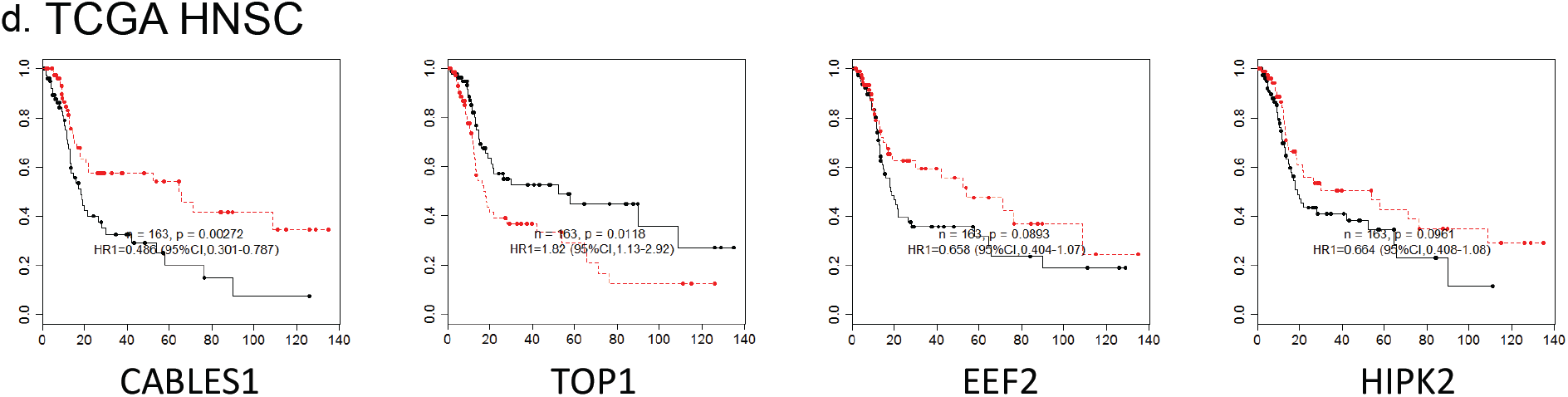
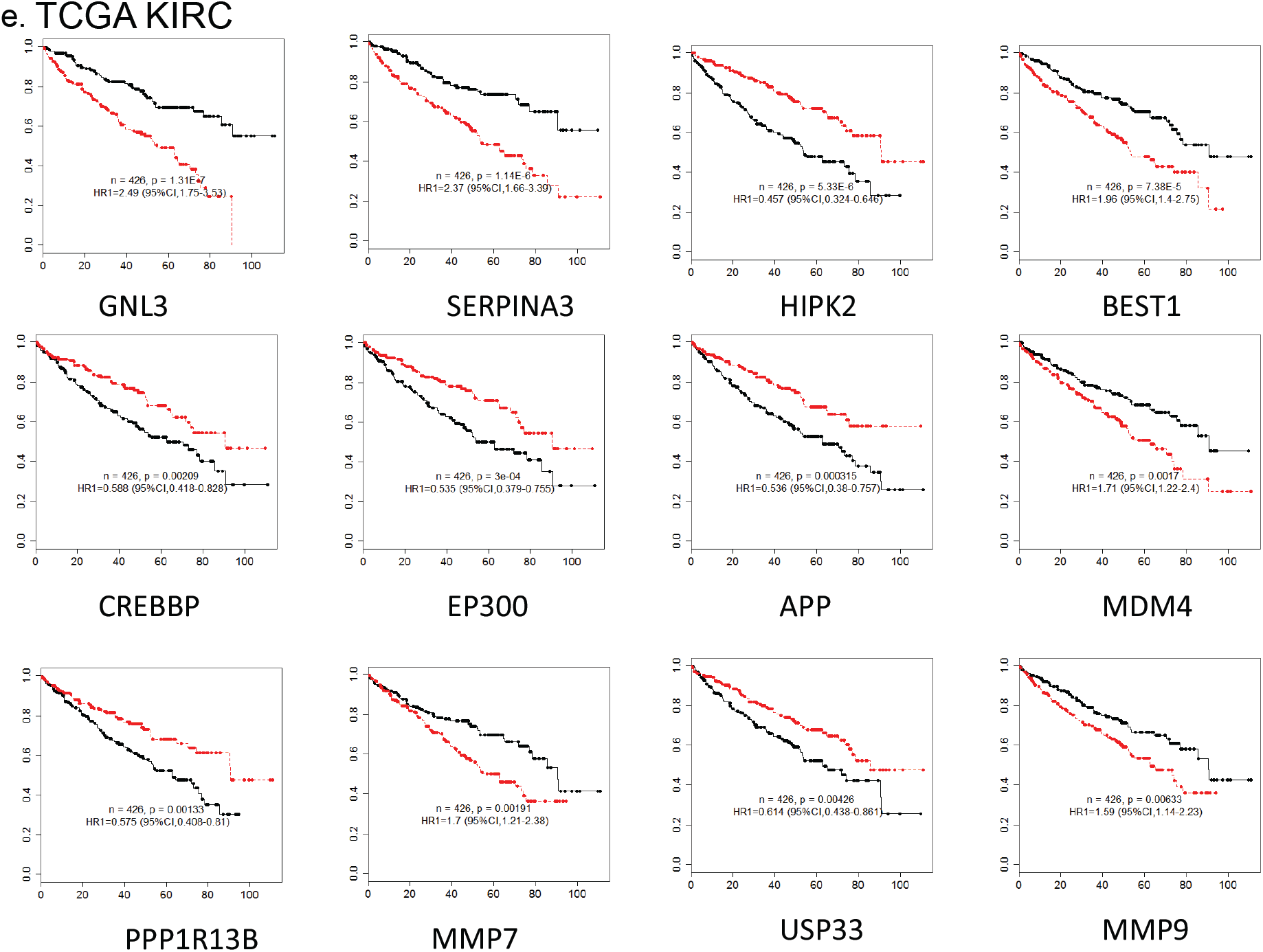
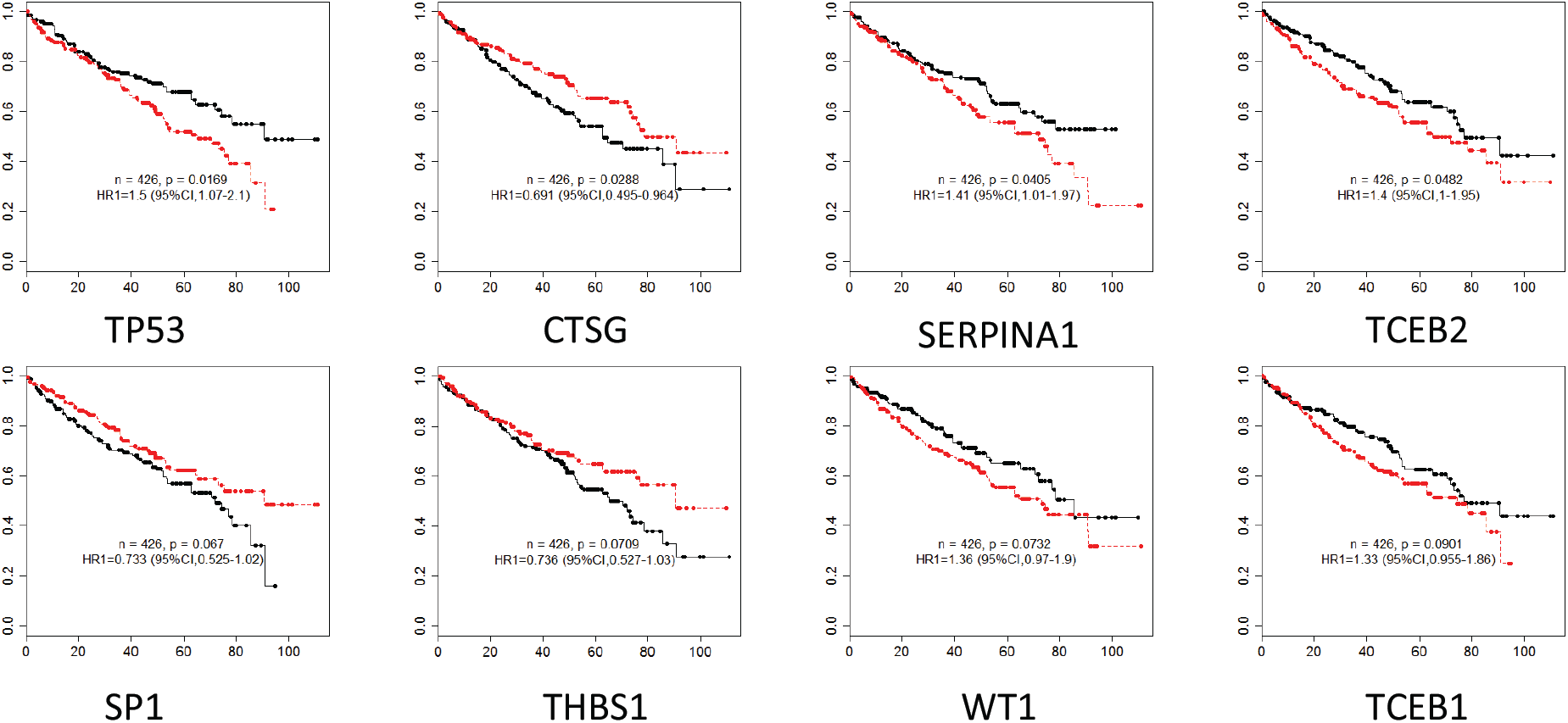
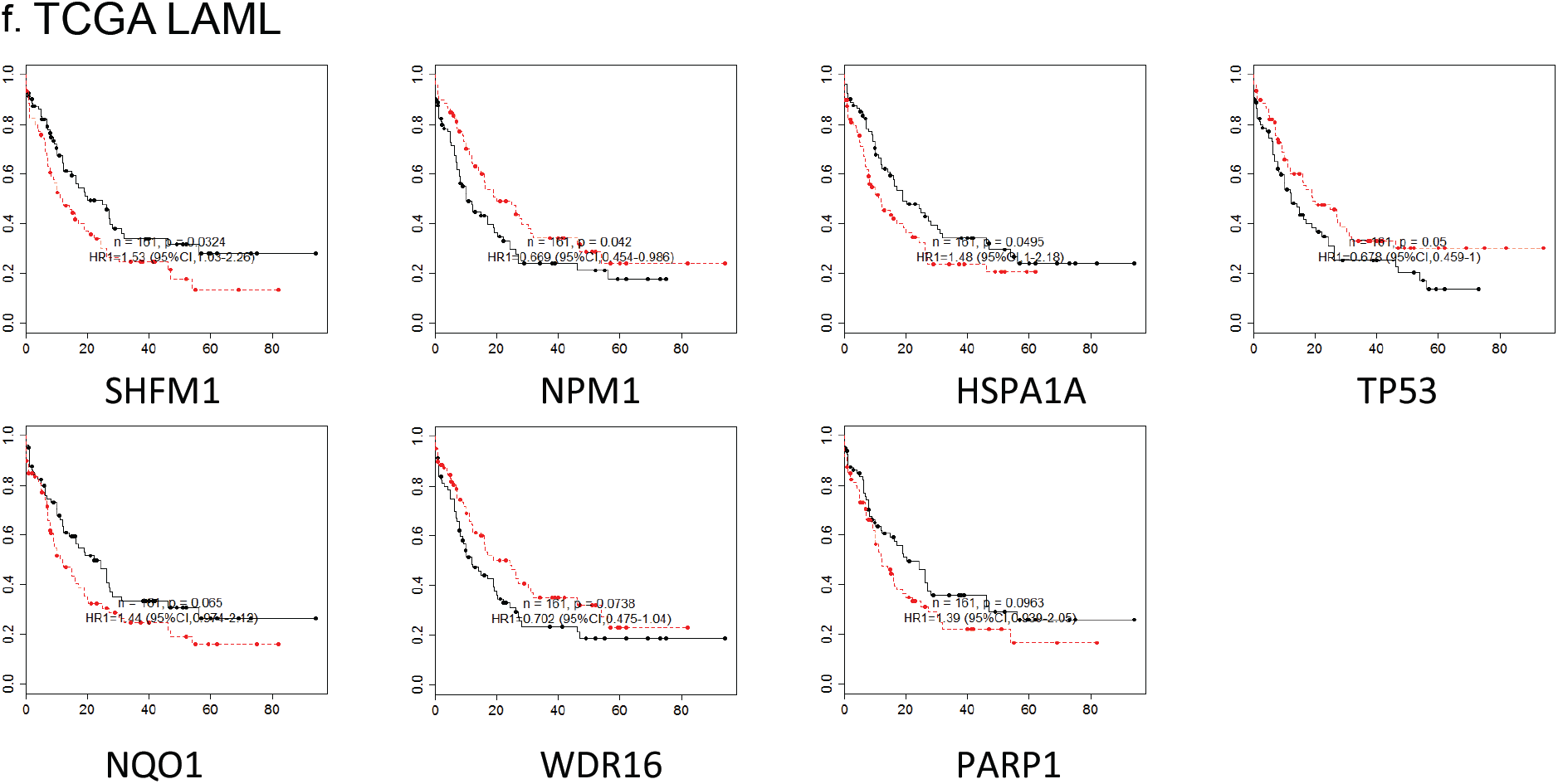
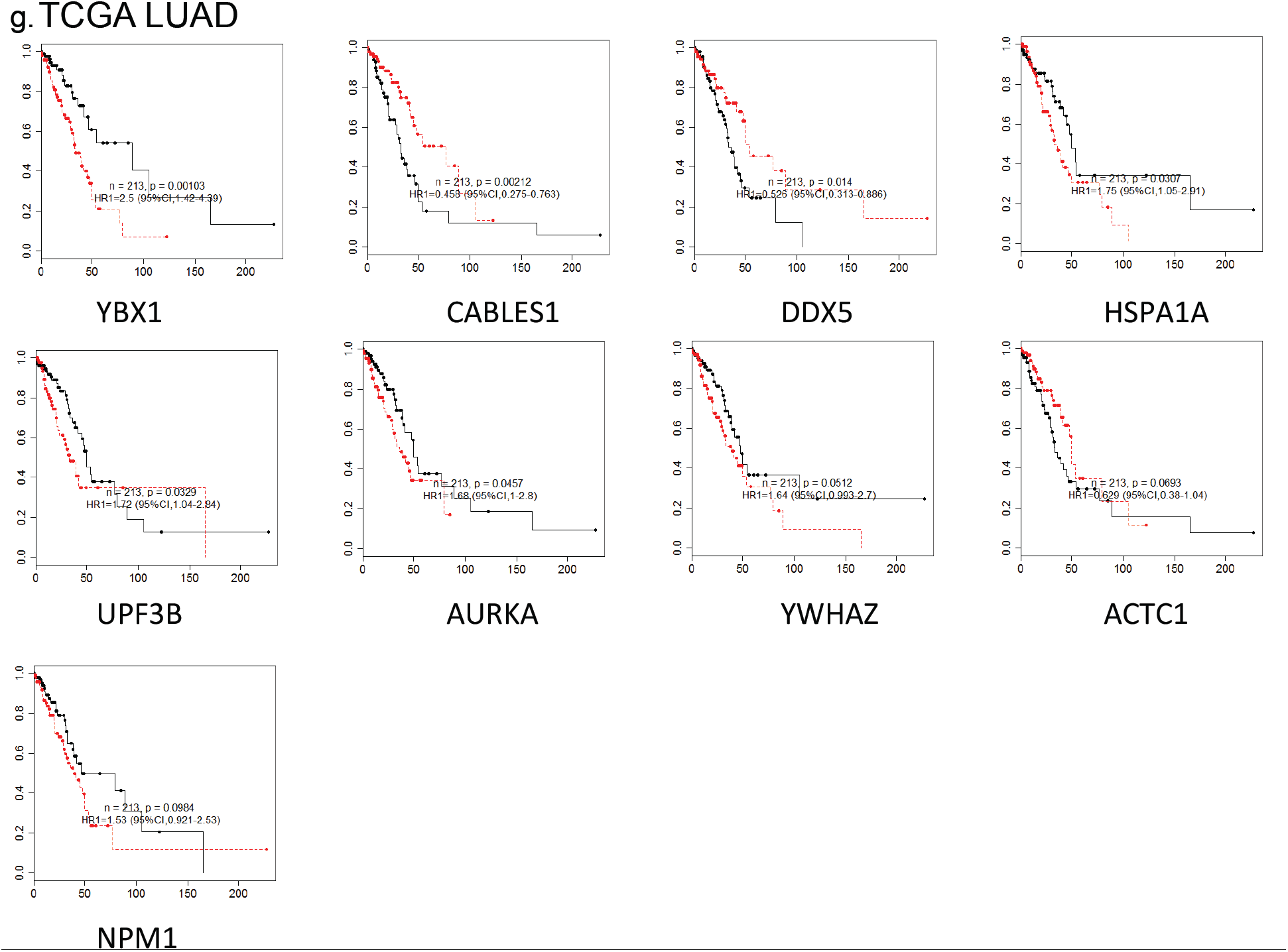
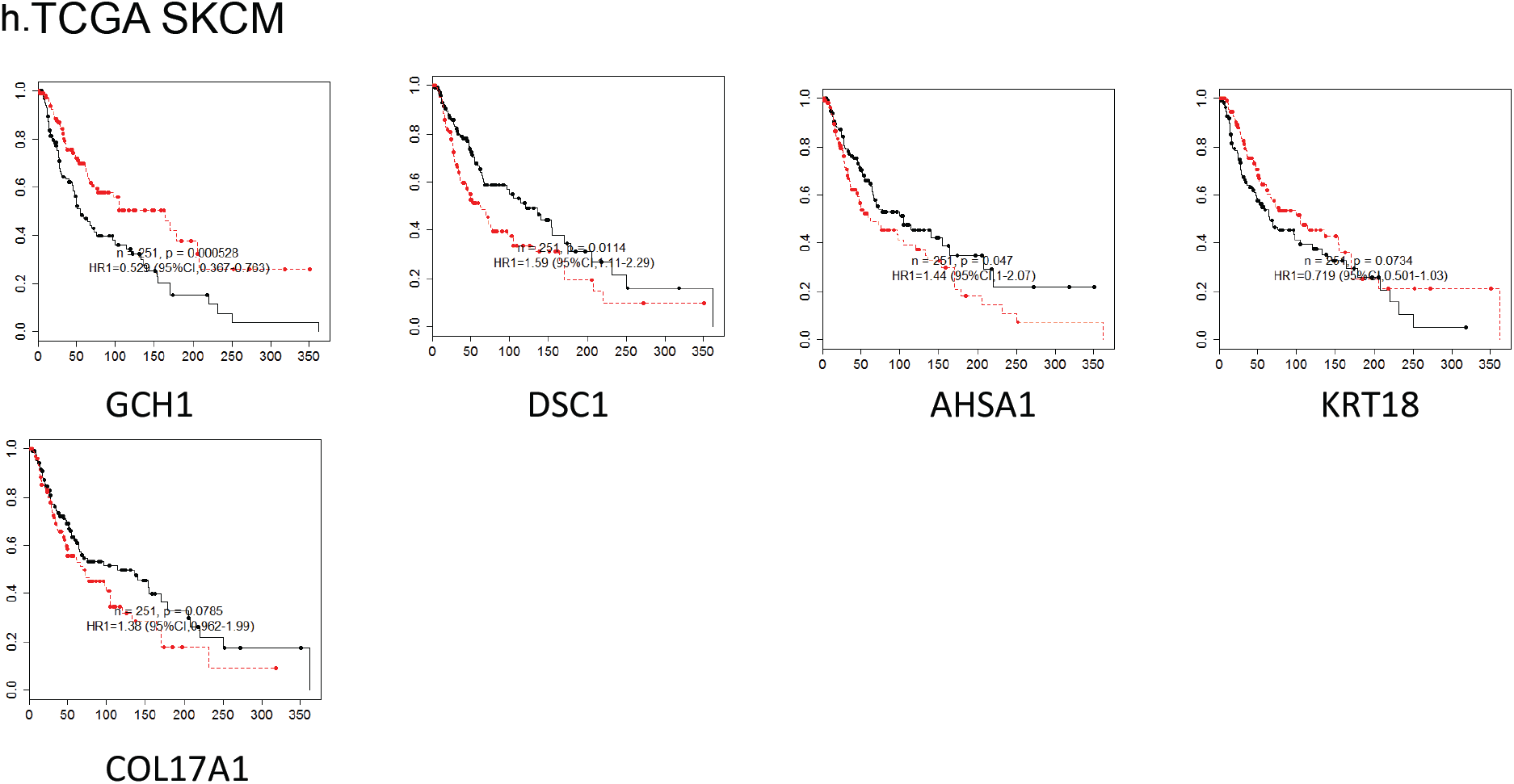
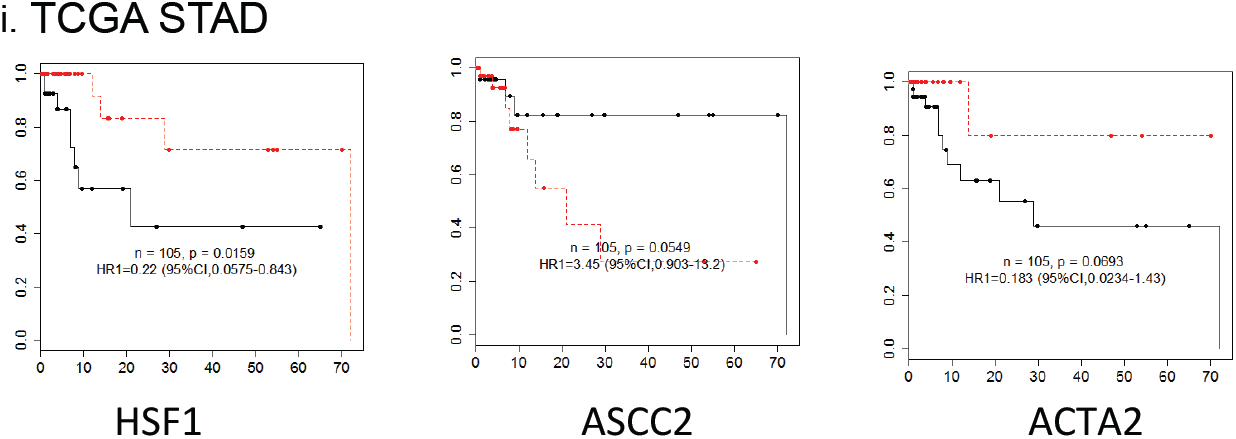
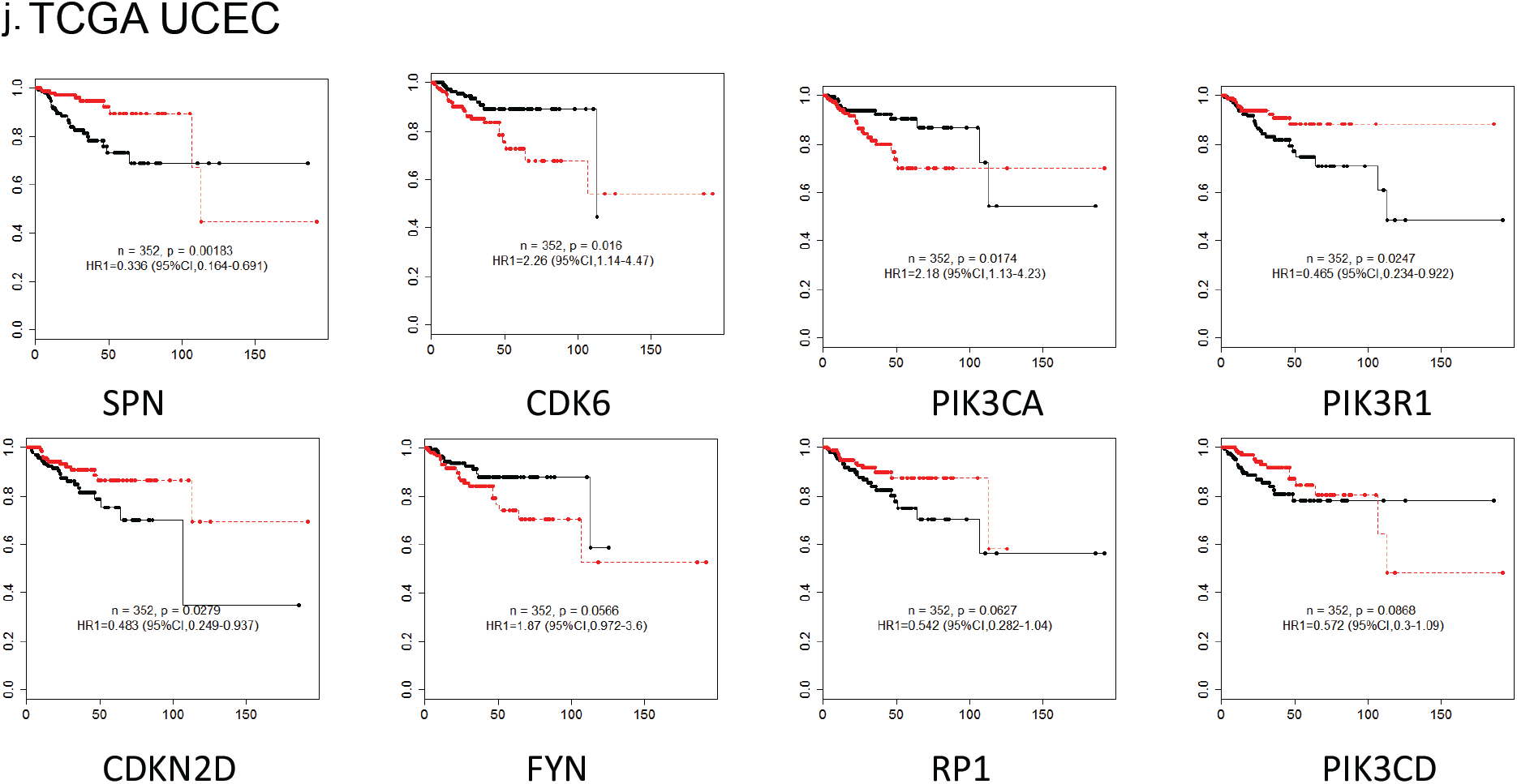
Supplementary Figure 5 Single gene analysis.

